# Population density triggers wing dimorphism via *NlNompC*-mediated insulin signaling in *Nilaparvata lugens*

**DOI:** 10.1101/2025.10.10.681384

**Authors:** Miao-Miao Tian, Yun-Fei Li, Zhuang-Zhuang Qiao, Xun-Kun Zhu, Dong Wen, Shu-Chang Wang, Wei-Hua Ma, Zhi-Hui Zhu, Hong-Xia Hua

## Abstract

The *Nilaparvata lugens* employs wing dimorphism as a key adaptive strategy to environmental heterogeneity. While hormonal pathways and transcription factors governing nutrient-induced wing plasticity are known, how population density triggers developmental divergence remains unresolved. Here, we mechanistically dissect density-dependent wing morph determination, revealing tactile inter-individual contact as the primary signal. High-density conditions amplify gentle-touch frequency, which is detected by mechanosensitive ion channels *Nl*NompC and *Nl*NMDARs. Genetic silencing or pharmacological inhibition of these channels abolished density-induced long-winged morphogenesis. The conserved role of *NlNompC* in gentle-touch sensation was validated by rescuing tactile perception deficits in *Drosophila nompC* mutants. Crucially, *Nl*NompC integrates tactile cues into insulin/insulin-like growth factor signaling (IIS): crowding-driven mechanical stimuli upregulate *NlIlp3*, while suppressing *NlInR2*, to activate wing elongation programs. Epistasis experiments confirmed IIS as the key downstream pathway, with co-silencing *NlInR2* or *NlFoxO* rescuing the short-winged phenotype induced by *NlNompC* knockdown or low population density, silencing *NlIlp3* or *NlInR1* rescuing the long-winged phenotype induced by high population density. Our work uncovers a mechano-endocrine axis linking tactile perception to developmental plasticity, bridging ecological cues with molecular pathways. These findings redefine density sensing beyond chemical modalities and offer actionable targets for disrupting pest dispersal strategies, advancing sustainable agriculture.

## Main

The brown planthopper (*Nilaparvata lugens*, BPH) represents a major pest species in rice cultivation systems, exhibiting wing dimorphism (long-winged versus short-winged morphs) regulated by multiple environmental factors including population density, host plant quality, etc (Kisimoto 1956, Saxena et al 1981, Iwanaga et al 1987). While existing research has elucidated the involvement of endocrine pathways (Iwanaga et al 1986, Xu et al 2015, Zhao et al 2017, Ye et al 2019) along with transcription factors such as *NlZfh1* (Zhang et al 2022) and wing development-related genes like *vestigial* (*Nlvg*) (Zhang et al 2021) and *ultrabithorax* (*Nlubx*) (Liu et al 2020) in nutrient-induced wing morph transition, critical knowledge gaps persist in three fundamental aspects in crowding-regulated wing dimorphism in BPH: (1) The precise signaling modality through which population density - a pivotal environmental determinant - regulates wing morph differentiation remains undefined; (2) The primary molecular mechanisms underlying crowding perception in BPH continue to be elusive; and (3) The signal transduction cascade mediating environmental cues to wing bud development for wing morph determination has yet to be fully elucidated. Understanding the nature of the stimuli associated with population density, whether they are visual, olfactory, or mechanical, and how BPH detects these signals, particularly the top-level sensory molecules involved in this process, is vital for a comprehensive understanding of its adaptive strategies.

### Mechanical Contact Mediates Density-Dependent Wing Dimorphism in BPH

To validate population density-dependent wing dimorphism, we conducted density-controlled rearing experiments in 17-mL plastic tubes containing rice seedlings. Female BPH exhibited pronounced density-dependent wing plasticity, with long-winged morph proportion increasing significantly at higher densities (Fig. 1a). In contrast, male wing morph remained unaffected, maintaining around 90% long-winged individuals across all densities (Fig. 1b), consistent with previous reports (Kisimoto 1956, Iwanaga et al 1985).

**Fig 1.**
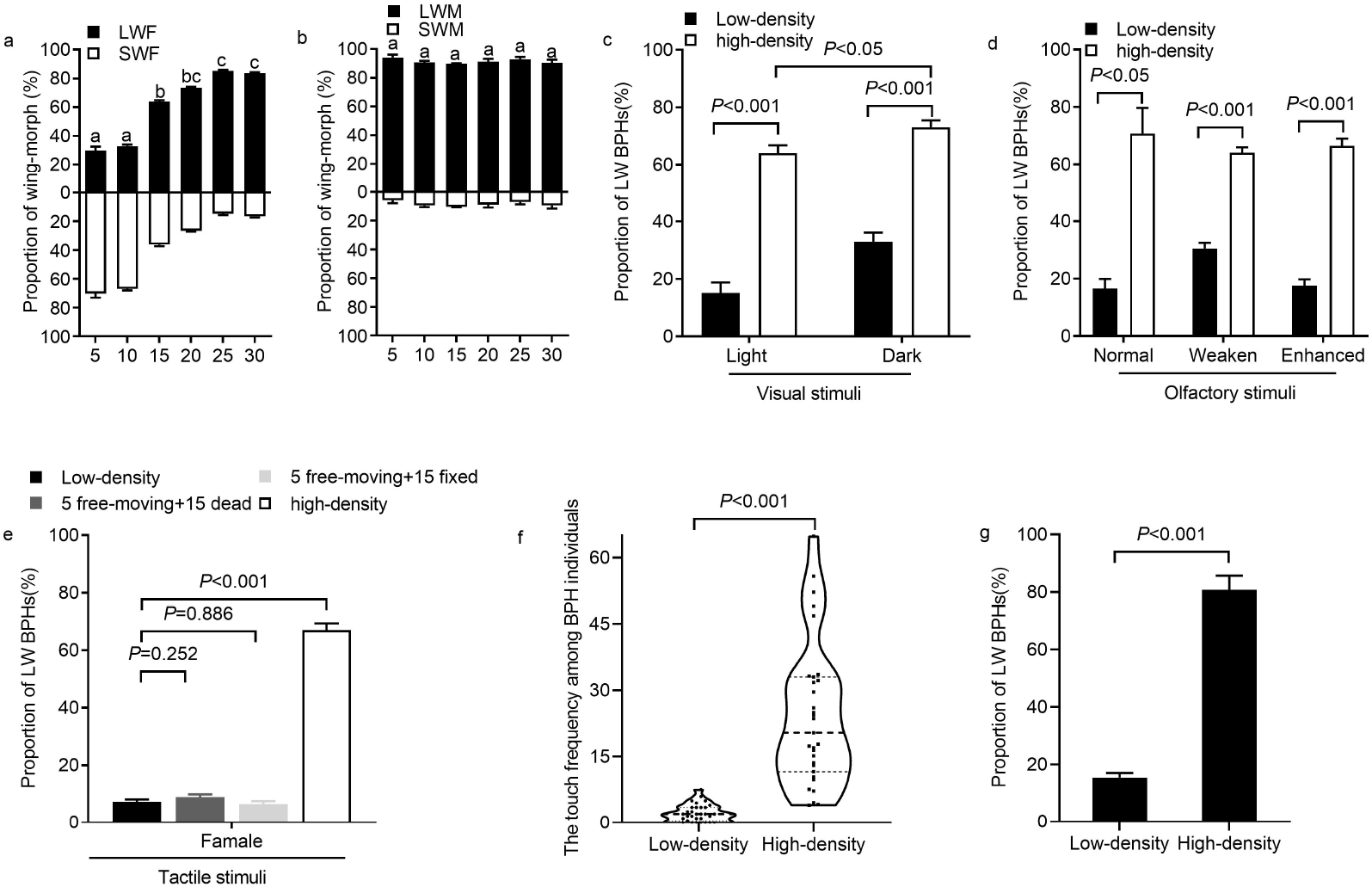
Crowding-induced long-winged morph formation of BPH is primarily mediated by tactile stimuli. **a-b**, 5, 10, 15, 20, 25 or 30 3rd nymphs of BPHs were reared in 17 ml vials, and wing morphs were counted after emergency. Proportion of wing morph of female (a) and male (b) BPHs. LWF: long-winged female, SWM: short-winged male. Means ± standard error of the mean (s.e. m). One-way analysis of variance using Duncan’s multiple range test, different letters indicate significant level at *P* < 0.05. n=40-57. **c-e**, The influence of various stimuli on wing morph ratio with high-density (20 nymphs) and low-density (5 nymphs) BPH treatments (in 17 ml vial), **c**, Visual stimuli, Light: low-density, or high-density BPHs reared in regular photoperiod (14L:10D) until adult emergence; Dark: low-density, or high-density BPHs reared in darkness until adult emergence; **d**, Olfactory stimuli: Weaken: antennae were amputated at the base of the third segment of antennae; Enhanced: 200 BPHs were kept outside (in 500 ml beaker) of the feeding vial; **e**, Tactile stimuli: 5 free-moving + 15 dead: 5 free-moving BPHs plus 15 dead BPHs, 5 free-moving + 15 fixed: 5 free-moving BPHs plus 15 live BPHs wrapped with gauze so that two groups could not touch. low-density (5 BPHs), or high-density (20 BPHs). **f and g**, BPHs were reared in a 3.5 ml container at low density of 2 nymphs and high density of 5 nymphs. While the absolute numbers differ between the 3.5 ml and 17 ml treatments (in a-c), the relative population density ratios (high/low) are comparable. n=15-60. **f**, The median gentile touch times among BPHs were calculated for (high-density (5), low-density (2) treaments (31 replications), as the data distribution was skewed. **g**, The wing-morph ratio of female in f. *P*<0.05 and *P*< 0.001 (Independent samples *t*-test). n=31.

Population density signals could involve visual, olfactory, or tactile cues. To identify the primary signal modality, we designed three experimental paradigms (Fig. S1). First, 24-hour dark conditions eliminated visual input but failed to reduce long-winged morph proportion in high-density groups (20 nymphs) compared to light-cycle controls, though low-density groups (5 nymphs) under darkness showed a slight increase in long-winged morphs (Fig. 1c). These results exclude visual cues as critical density signals.

Next, we disrupted olfactory signaling by either ablating antennal flagella (reducing pheromone reception, Weaken) or surrounding test tubes with 200 additional BPHs (equalizing pheromone gradients across densities, Enhanced). Neither intervention altered density-dependent wing differentiation (Fig. 1d), ruling out olfactory cues as primary mediators.

Direct mechanical stimulation assays used in aphids (Johnson et al 1965) and in locusts (Simpson et al 2001) proved infeasible due to BPH’s small size and hyperactivity. Instead, we indirectly tested tactile signaling by adding 15 dead nymphs (5 free-moving + 15 dead) or 15 live nymphs physically separated by gauze (5 free-moving + 15 fixed) to low-density groups (5 nymphs). Both treatments failed to mimic high-density effects, with long-winged frequencies remaining indistinguishable from unmanipulated low-density groups and significantly lower than high-density controls (20 nymphs) (Fig. 1e). Spatial segregation thus abolished crowding-dependent wing induction, implicating direct inter-individual contact as essential.

To quantify tactile interactions, we video-recorded 3rd-instar nymphs in 3.5-mL cuvettes under a JH-T6 microscope for 24 hours. Even at low density (2 nymphs), sporadic contacts occurred, but high density (5 nymphs) significantly increased gentle touch frequency (Fig. 1f), correlating positively with long-winged morph proportions (Fig. 1g).

Collectively, these results establish inter-individual gentle touch as the principal density signal driving wing dimorphism in BPH, with minimal contributions from visual or olfactory cues. This parallels aphid wing plasticity under crowding (Johnson et al 1965), highlighting conserved mechanosensory regulation of developmental polyphenism.

### Mechanosensitive Ion Channels Mediate Crowding-Induced Wing Morph Differentiation via Gentle Tactile Sensing

In *Drosophila*, gentle-touch sensation is mediated by ion channels including No mechanoreceptor potential C (NompC), heteromeric N-methyl-D-aspartate receptors (NMDAR1/NMDAR2), and Pickpocket (ppk) (Turner et al 2016). However, BLAST analysis of the BPH genome identified only orthologs of NompC, NMDAR1, and NMDAR2. To pinpoint the mechanotransducers governing wing dimorphism, we performed RNAi-mediated knockdown of these candidates under high-density rearing (20 nymphs per tube). Silencing *NlNompC* reduced long-winged morph proportion in females from 61% to 24% and in males from 70% to 37% (Fig S10a; Figs. 2a, b). Similar suppression occurred with a second *NlNompC*-targeting dsRNA (ds2*NlNompC*; Figs. S2a, b). Knockdown of *NlNMDAR1* or *NlNMDAR2* also significantly reduced long-winged morph proportions in both sexes (Fig. 2c; Figs. S2c–e; Fig S10 b, c).

**Fig 2.**
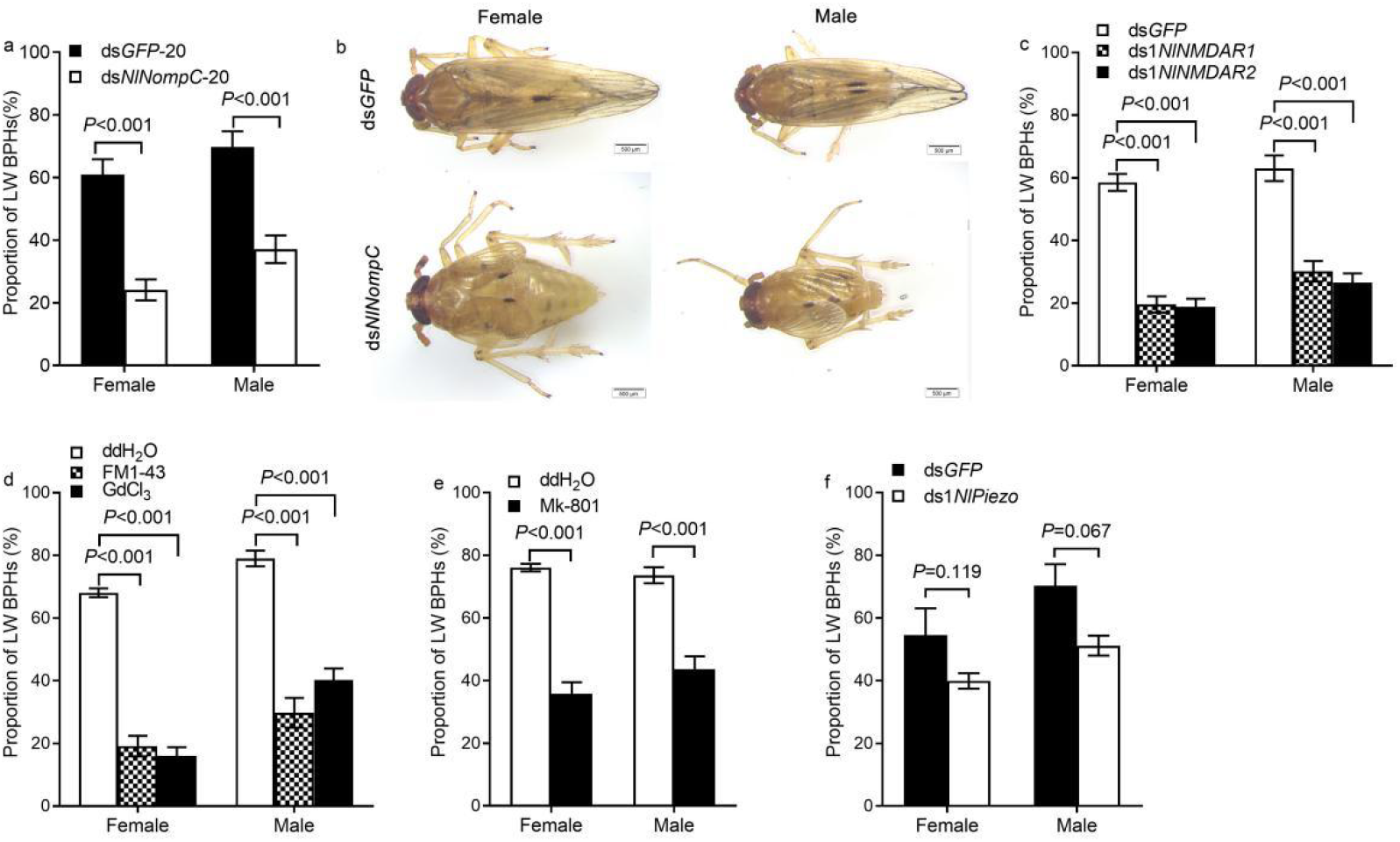
Mechanosensitive ion channels for gentle-touch sensation mediate crowding-induced wing morph differentiation. **a**, The wing morph ratio of BPHs injected with ds*GFP* or ds*NlNompC* and reared in 17 ml vial with density of 20 nymphal BPHs (for ds*GFP*-20, ds*NlNompC*-20) **b**, Treatments with ds*GFP*, ds*NlNompC*. **c-f**, The wing morph ratio of BPHs injected with ds*GFP* or ds*NlNMDAR1/NlNMDAR2* (c) ddH_2_O or FM1-43/GdCl_3_ (d), ddH_2_O or Mk-801 (e) and ds*GFP* or ds*NlPiezo* (f) in 17 ml vial with density of 20 nymphal BPHs. Independent samples *t*-test, n=12-58.

As *NlNompC* is essential and thus refractory to knockout, we pharmacologically inhibited its activity. Injecting GdCl_3_ (6 × 10^−12^ µmol/nymph) or FM1-43 (1.2 × 10^−12^ µmol/nymph) suppressed long-winged morph differentiation (Fig. 2d) (Yan et al 2013). Similarly, injection of NMDAR antagonist MK-801 (6 × 10^−12^ µmol/nymph) abolished density-dependent wing plasticity (Fig. 2e) (Wong et al 1986; Geister et al 2008) .

Insect mechanosensation also involves nociceptive channels like Piezo and Painless, which detect harmful stimuli (Tracey et al 2003, Kim et al 2012). To test if nociception contributes to wing morphogenesis, we silenced *NlPiezo*. Unlike *NlNompC*/*NMDAR*s knockdown, *NlPiezo* RNAi had no significant effect on long-winged morph proportion (Fig. 2f; Figs. S2f, g; Fig S10d), unequivocally implicating gentle-touch-sensitive channels—not nociceptive pathways—in density-induced wing differentiation. This also aligns with the behavioral impacts of elevated population density in BPH: increased locomotion and mortality under crowding, yet no escalation in harmful interactions such as fighting or cannibalism among individuals.

### *Nl*NompC Senses Gentle Tactile Cues to Regulate Wing Dimorphism in BPH

In *Drosophila*, NompC mediates gentle-touch sensation and modulates locomotion (Cheng et al 2010, Yan et al 2013). BPH *Nl*NompC shares 83.13% amino acid identity with its *Drosophila* counterpart, including 29 conserved microtubule-associated ankyrin repeats (Fig. S3). Fluorescence in situ hybridization (FISH) revealed high *NlNompC* expression in the epidermis of leg, head, thorax and abdomen (Fig. 3a; Fig. S4). RNAi-mediated *NlNompC* knockdown significantly impaired BPH nymphs’ gentle-touch responsiveness (Wang et al 2019), preliminarily confirming its mechanosensory role, as observed in its *Drosophila* ortholog NompC. To further validate *Nl*NompC’s tactile sensing function, we expressed *Nl*NompC in in class III dendritic arborization neurons of *Drosophila nompC* mutants (genotype: *19-12-Gal4* > *UAS*-*NlNompC*; *nompC*^f00914/nompC1^) (Yan et al 2013, Li et al 2016). This rescued the tactile perception deficits in *nompC* mutants (Fig. S5; Fig. 3b, c), further demonstrating functional conservation of *Nl*NompC as a gentle-touch sensor.

**Fig 3.**
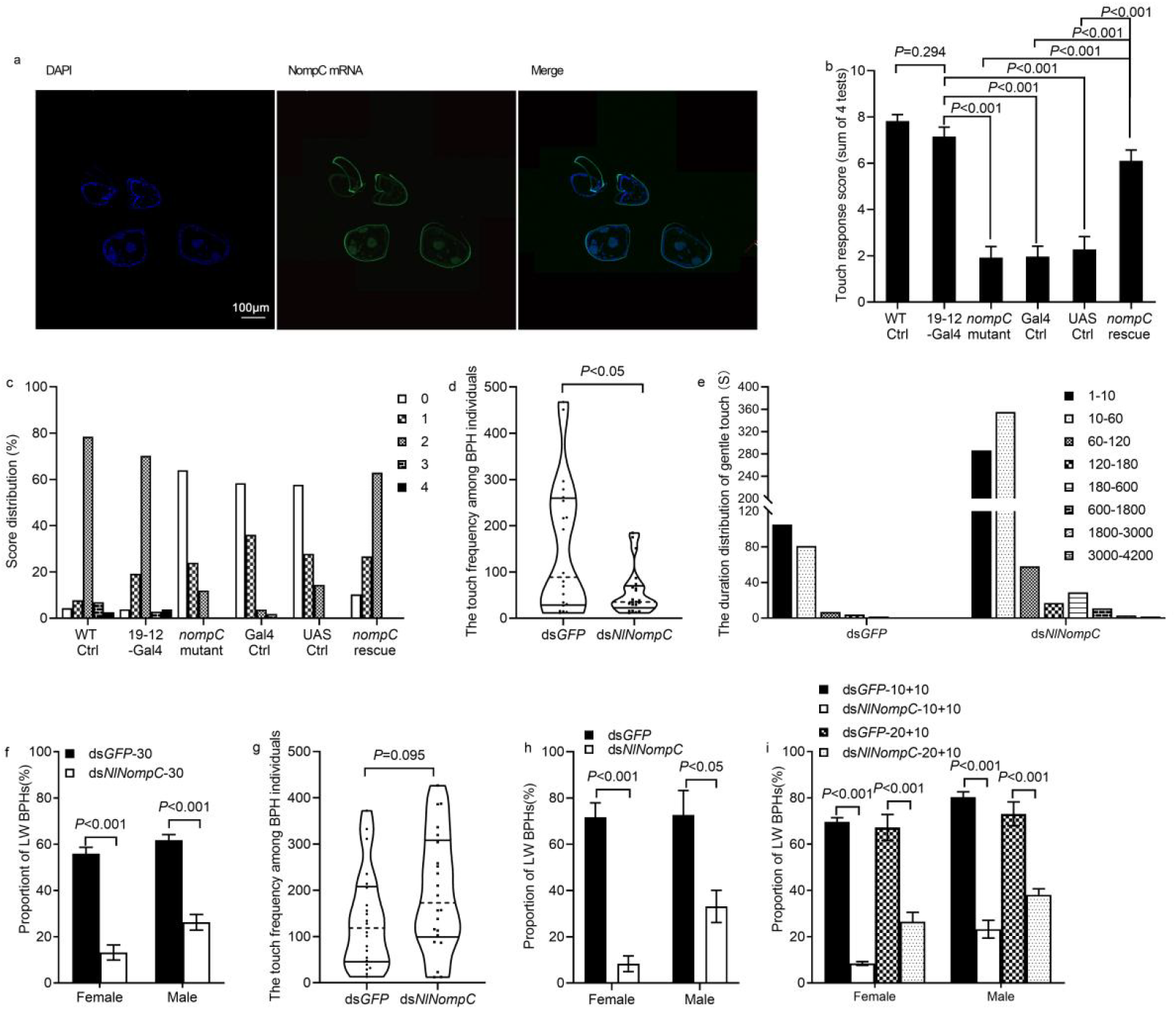
*NlNompC* functions as a gentle-touch receptor. **a**, Localization of *NlNompC* mRNA in legs. The oligonucleotide probe labeled with fluorochrome FAM (5’ primer sense) targeting *NlNompC* expressed in epidermis. Scale bars: 100 μm. **b-c**. Defects of gentle-touch sensation in larvae of *Drosophila nompC* mutant *nompC*^*f00914*^ */nompC*^*1*^ could be significantly rescued by expressing *Nl*NompC in class III dendritic arborization neurons (*nompC* rescue), but not by UAS-*Nl*NompC (UAS Ctrl) or Gal4 (Gal4 Ctrl) alone. *nompC* mutant, Gal4 Ctrl and UAS Ctrl were compared to both 19-12-Gal4 and *nompC* rescue for the significance test. **b**, Touch response score of different genotypes. **c**, Score distribution of different genotypes. Genotypes are as follows: 19-12-Gal4: *repo-GAL80*/+; *19-12-GAL4*/+. *nompC* mutant: *nompC*^*f00914*^*/nompC*^*1*^; +/+. Gal4 control: *nompC*^*f00914*^*/nompC*^*1*^; *19-12-GAL4*/+. UAS control: *nompC*^*f00914*^*/nompC*^*1*^; *UAS-NlnompC*/+. *nompC* rescue: *nompC*^*f00914*^*/nompC*^*1*^; *UAS-NlnompC*/ *19-12-GAL4*. Wild-type control: *w*^*1118*^. (n =24–29). **d-e**, dsRNA injected BPHs were reared in 3.5 ml container with density of 5 nymphs (n=20-22). **d**. The median gentle touch frequency among planthoppers treated with ds*GFP*, ds*NlNompC*. **e**. The duration (seconds) of each gentile touch of planthoppers treated with ds*GFP*, ds*NlNompC. P*<0.05 and *P*< 0.001 (Independent samples *t*-test). **f**, The wing morph ratio of BPHs injected with ds*GFP* or ds*NlNompC* and reared in 17 ml vial with density of 30 nymphal BPHs (ds*GFP*-30 and ds*NlNompC*-30, respectively, n=39-51). **g-h**, dsRNA injected BPHs were reared in 3.5 ml container with density of 5 nymphs plus 2 uninjected nymphs of *S. furcifera*. (n=20-22) **g**, The median gentile touch frequency among planthoppers. **h**, The wing-morph ratio of BPH. **i**, The wing morph ratio of BPHs injected with ds*GFP* or ds*NlNompC* and reared in 17 ml vial with density 10 or 20 dsRNA injected BPHs plus 10 unjected nymphs of *S. furcifera* (for ds*GFP*-10+10, ds*NlNompC*-10+10, ds*GFP*-20+10, ds*NlNompC*-20+10, respectively, right 2 panels). *P*<0.05 and *P*< 0.001 (Independent samples *t*-test, n=55-56)

Notably, *NlNompC* silencing also reduced nymphal motility, decreasing inter-individual contact frequency from 89 contacts/24 h in ds*GFP* controls to 35 contacts/24 h (Fig. 3d, Fig. S2a). However, prolonged contact durations were observed in ds*NlNompC* nymphs (max 3,847 sec vs. 153 sec in controls), often involving one nymph climbing and remaining atop another (Fig. 3e; Movies S1-S4). This aberrant behavior suggests *Nl*NompC loss disrupts sensation of and therefore defensive responses to tactile stimuli among BPHs.

To disentangle whether wing morph defects in *NlNompC*-silenced nymphs stem from reduced locomotion (lower contact frequency) or impaired mechanosensation, we conducted two compensatory experiments: 1. Density compensation: Raising rearing density to 30 nymphs/tube (vs. 20 nymphs in standard high-density conditions) failed to restore long-winged morph frequencies in *NlNompC* knockdown groups (female: 13% vs. 55% in ds*GFP*; male: 26% vs. 60%; Fig. 3f; Fig S10 e). 2. Interspecific contact supplementation: Adding closely related *Sogatella furcifera* nymphs restored inter-individual contact frequencies among planthoppers in *NlNompC*-silenced BPH (Fig. 3g; Fig S10 f-g), yet inter-individual contact duration remained significantly prolonged (Fig. S6 b, Movies S5-S8). However, the long-winged morph proportion remained reduced (Figs. 3h, i; Fig S10 f-g). These findings collectively indicate that *Nl*NompC’s mechanosensory role is essential for density-dependent wing polyphenism in BPH, as its knockdown disrupts long-winged morph development even under conditions of unaltered gentle-touch signal input. We modeled the cumulative tactile stimulus input (S_24_) in 24 h as: S_24_=S_st_×F×D, where S_st_ = single-touch stimulus strength, F = contact frequency, and D = contact duration. Despite reduced F in *NlNompC* knockdown nymphs, prolonged D (Fig. 3e; Fig. S6b; Movies S1-S8) likely preserves S_24_, yet long-winged morphogenesis remains disrupted (Fig. S6 a). This further confirms that *Nl*NompC-dependent mechanosensation is indispensable for translating tactile cues into developmental outcomes.

### *Nl*NompC Regulates Wing Dimorphism via Insulin/IGF Signaling (IIS) Pathway

To elucidate how mechanosensory inputs are integrated into wing developmental programming, we investigated hormonal pathways implicated in insect polyphenism. Quantitative PCR revealed no significant expression changes in juvenile hormone-related genes *Krüppel homolog 1* (*NlKr-h1*) and *Methoprene-tolerant* (*NlMet*) or ecdysteroid receptor *ultraspiracle protein* (*NlUSP*) under *NlNompC* knockdown or density variation (Jindra et al 2013, Yamanaka et al 2013 ). While ecdysteroid-responsive genes *Ecdysteroid receptor* (*NlEcR*) and *Ecdysone-induced protein 75* (*NlE75*) were suppressed by *NlNompC* silencing, their expression exhibited inverse trends across density gradients (Figs. S7a, b). These results exclude JH and ecdysteroid signaling as central regulators of crowding-induced wing morphogenesis, pivoting focus to insulin signaling as the primary regulatory axis.

Focusing on insulin signaling, we observed IIS pathway activation correlated with mechanosensation. *NlNompC* silencing under high density significantly downregulated *NlIlp3* (*insulin-like peptide 3*), *NlAkt* (*kinase B*), and wing determinant *Nlvg*, while upregulating *NlInR2* (*insulin receptor 2*) (Fig. 4a) (Xu et al 2015). Similarly, low-density conditions suppressed *NlIlp3* and *NlInR1* but elevated *NlInR2* (Fig. 4b). Western blot and immunofluorescence confirmed reduced *Nl*Ilp3 protein levels in *NlNompC*-silenced (Figs. 4c, d; Fig. S8) or low-density nymphs (Figs. 4e, f), establishing IIS as a downstream mediator of population density and mechanosensory input.

**Fig 4.**
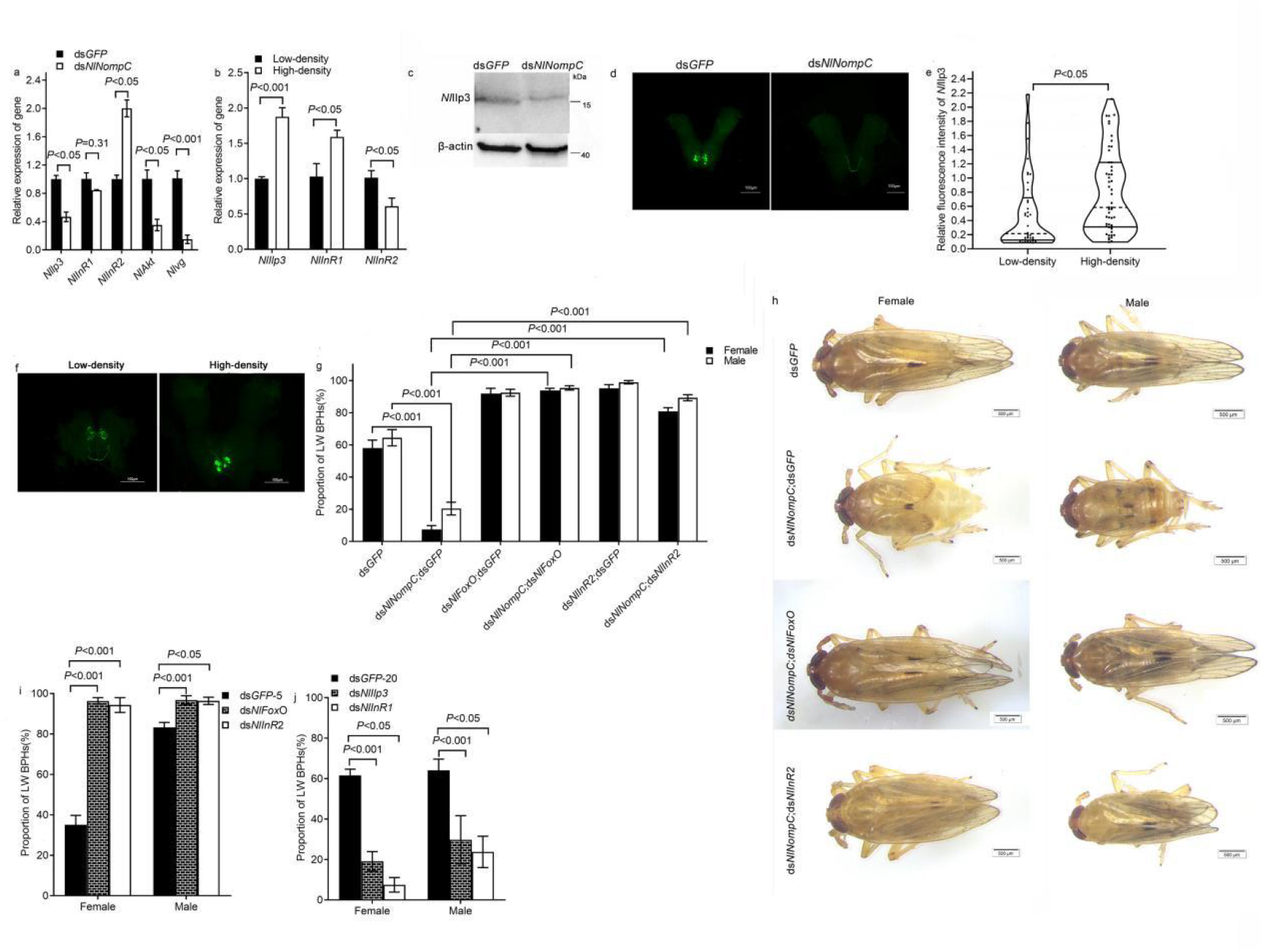
*Nl*NompC mediated wing morph switch of BPH is relayed by the insulin signaling. **a**, The expression level of *NlIlp3, NlInR1, NlInR2, NlIAkt* and *Nlvg* of 5th-instar nymphs after treated with ds*GFP* or ds*NlNompC*. **b**, The expression level of *NlIlp3, NlInR1, NlInR2* of 5th-instar nymphs in low-density (5 nymphs/tube) and high-density (20 nymphs/tube). **c**. The *Nl*Ilp3 level in brains was assessed by Western blotting (WB) with polyclone antibody raised by injection of rabbit with the first 118 amino acids of *Nl*Ilp3 protein produced in *E. coli* after *treated* with ds*GFP* or ds*NlNompC*. **c**, Immunofluorescence detection of *Nl*Ilp3 in brains of BPHs after *treated* with ds*GFP* or ds*NlNompC*. n=20-52. **e**, Relative fluorescence intensity of *Nl*Ilp3 of 5th-instar nymphs in low-density and high-density. n=38-51. **f**. Immunofluorescence detection of *Nl*Ilp3 of 5th-instar nymphs in low-density and high-density. n=38-51. **g**, ds*NlFoxO* or ds*NlInR2* reverses ds*NlNompC* effect (n=11-33). **h**, Treatments with ds*GFP*, ds*NlNompC*;ds*GFP*, ds*NlNompC*;ds*NlFoxO* or ds*NlNompC*;ds*NlInR2*. **i**,Treatments with ds*GFP*, ds*NlFoxO* or ds*NlInR2* in low-density rearing condition. n=24-43. **j**,Treatments with ds*GFP*, ds*NlIlp3* or ds*NlInR1* in high-density rearing condition. n=11-14. *P*<0.05 and *P*< 0.001 (Independent samples *t*-test)

**Fig 5.**
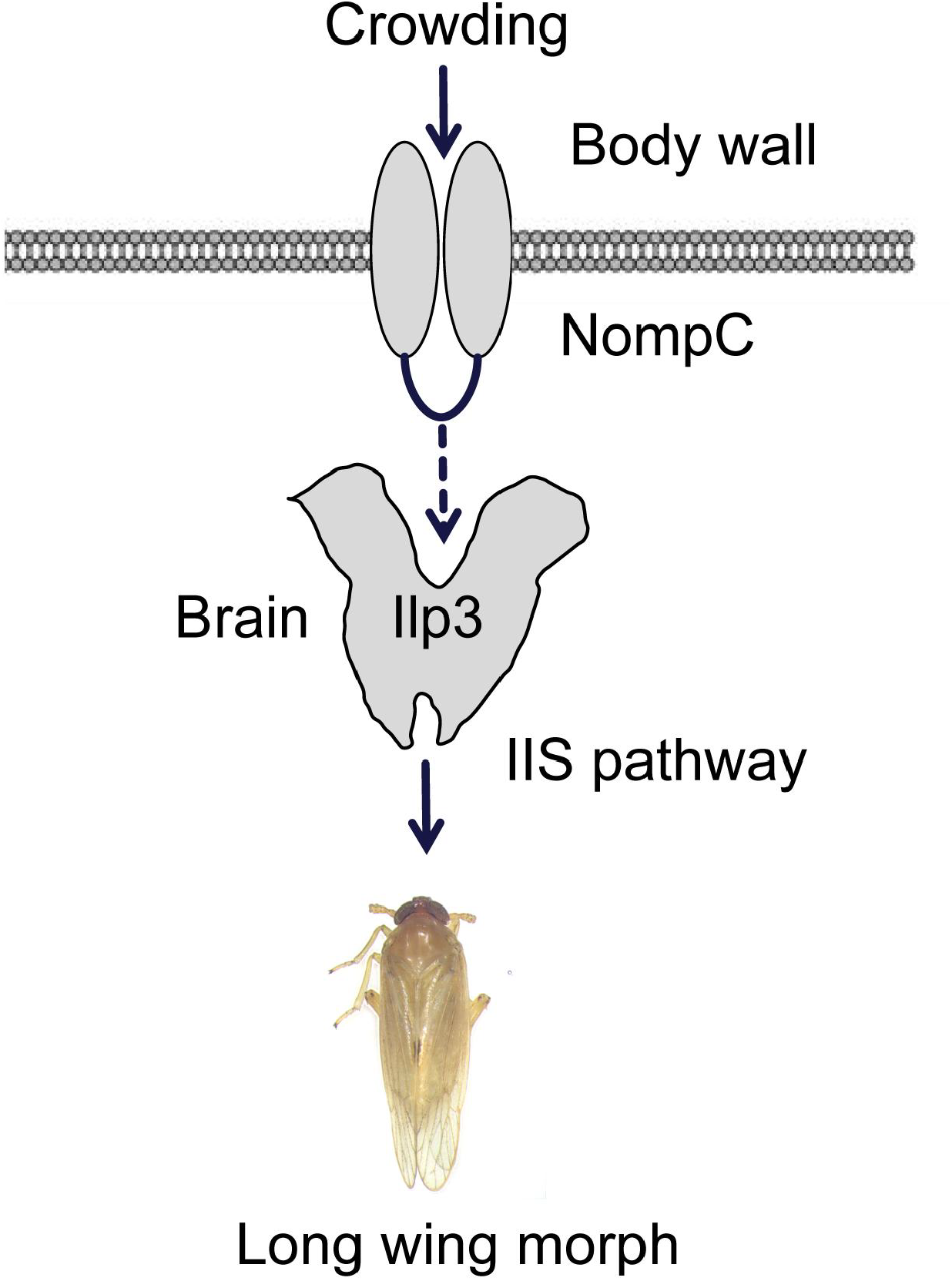
Model of the molecular regulation of wing polyphenism in population density of BPHs. Elevated population density increases inter-individual tactile contact, which serves as the primary density signal. This gentle tactile stimulation is perceived by mechanosensitive ion channels, including *NlNompC* and *NlNMDARs*, and transmitted via sensory neurons to the central nervous system (CNS). The CNS integrates these signals and directs insulin-producing cells (IPCs) to synthesize and secrete insulin-like peptide Ilp3. Subsequently, Ilp3 activates the insulin signaling pathway, which interfaces with the genetic network governing wing development, ultimately driving the transition to the long-winged morph.

To further validate this model, we performed genetic epistasis experiments by co-silencing *NlNompC* with *NlInR2* or *NlFoxO*—two short-wing-promoting genes (Figs. S10 h-n). Remarkably, over 80% of ds*NlNompC*;ds*NlInR2*- or ds*NlNompC*;ds*NlFoxO*-treated BPHs developed long-winged morphs, regardless of sex (Figs. 4g, h; Fig S10 h-n). Similarly, *NlInR2* or *NlFoxO* knockdown under low-density conditions (Figs. S10 o-p) induced near-complete long-winged differentiation (Fig. 4i). Conversely, silencing *NlIlp3* or *NlInR1* under high-density crowding (Figs. S10 q-r) suppressed long-winged morphs to <30% (Fig. 4j). These genetic manipulations conclusively demonstrate that mechanosensitive ion channels transduce population density signals via the insulin pathway to regulate wing dimorphism in *N. lugens*.

*NlNompC* silencing suppresses insulin signaling activity, as evidenced by reduced *NlIlp3* expression (Figs, 4a, c, d), which is predicted to impair organ growth (Texada et al 2020). Morphometric analyses confirmed this phenotype: *NlNompC* knockdown significantly decreased forewing area (Fig. S9a) and hind tibia length (Fig. S9b) in short-winged morphs. Strikingly, co-silencing *NlFoxO* or *NlInR2* with *NlNompC* rescued these defects, partially restoring forewing size (Fig. S9c) and fully restoring tibia length to near-wild-type levels (Fig. S9d). These findings genetically demonstrate that *Nl*NompC-dependent tactile sensing governs IIS activity to orchestrate developmental plasticity in *N. lugens*.

## Discussion

Insect wing dimorphism represents a pivotal survival strategy shaped by environmental cues, yet the molecular mechanisms underlying density perception and signal transduction remain poorly understood. Our study bridges this critical knowledge gap by elucidating how the BPH translates tactile stimuli from crowding into developmental decisions through mechanosensitive ion channels for gentle-touch sensation and insulin signaling.

Three key findings emerge. First, we identified gentle tactile contact, rather than visual or olfactory cues, as the primary density signal (Fig. 1). High-density conditions amplified inter-individual touch frequency, triggering long-winged morph differentiation in BPH nymphs. This parallels aphid mechanosensation (Johnson et al 1965) but contrasts with locusts’ multimodal density sensing (Guo et al 2020). Second, we demonstrated *Nl*NompC and *Nl*NMDARs, but not *Nl*Piezo as essential mechanotransducers through RNAi and pharmacological inhibition (Fig. 2). The functional conservation of *Nl*NompC was validated by rescuing touch sensitivity in *Drosophila nompC* mutants (Fig. 3b, c) and the defects of defensive responses among the BPH nymphs (Fig 3 e, Fig S6 b). Notably, while NompC’s role in locomotion complicates interpretation, our density compensation experiments and *S. furcifera* addition assays (Fig 3. f, i) conclusively established its sensory function in wing morph determination. In addition, microtubules play a crucial role in the transmission of mechanical stimuli (Zhang et al 2015, Yan et al 2018). Paclitaxel, a microtubule inhibitor that suppresses microtubule dynamics, reduced the proportion of long-winged morphs in BPH (Lin et al 2020), which further illustrates the role of mechanical stimuli and mechanosensitive ion channels in wing polyphenism regulation. Third, we mapped the signaling cascade from mechanical stimuli to insulin pathway activation. Mechanical inputs upregulated *Nl*Ilp3 expression (Fig. 4 b, e, f) while *NlNompC* knockdown suppressed IIS components (Fig. 4 a, c, d), with genetic epistasis confirming IIS as the ultimate wing determinant (Fig. 4 g-i).

While mechanical signaling through ion channels like Piezo1 is known to regulate local tissue development via direct mechanotransduction (such as vascular morphogenesis in mice) (Li et al 2014, Ranade et al 2014), our work uncovers a novel paradigm where tactile stimuli sensed by epidermis *Nl*NompC/*Nl*NMDARs are transduced into remote endocrine responses via the nervous system. This neural-endocrine relay mechanism expands the functional repertoire of mechanosensitive channels beyond local tissue patterning to systemic developmental regulation. The *Nl*NompC-IIS axis likely represents an evolutionarily conserved mechanism, given IIS’s conserved role in insect polyphenisms.

Population density profoundly influences animal behavior, reproduction, and developmental plasticity (Knell 2009, Edwards et al 2021, Rajendiran et al 2021, Guo et al 2020; Yang et al 2023). In species ranging from teleosts to locusts, density-dependent responses often involve chemical cues such as pheromones (such as 4-VA in locusts) ( Guo et al 2020) or environmental metabolites (such as oxygen levels in fish sex determination, Rajendiran et al 2021). Our findings provide a universal framework for understanding density-dependent adaptations across taxa. The identification of tactile sensing as a primary density signal in BPH challenges the prevailing chemical-centric view of density regulation and suggests mechanical perception may underpin unsolved cases of developmental plasticity in other species. For instance, aggression escalation in crowded vertebrates or reproductive suppression in voles could involve analogous mechanosensitive pathways yet to be characterized (Knell 2009, Edwards et al 2021).

Key questions remain unresolved: (1) How mechanical signals are transduced from ion channels to insulin-producing cells; (2) Mechanical stimulation signal accumulating counter induced by the duration of high population density; (3) Potential crosstalk between *Nl*NompC and other density-sensing modalities; (4) The basis for sexual dimorphism in density responses. Addressing these through single-cell transcriptomics, methylated genomics and in vivo calcium imaging could unveil neural circuit mechanisms. Furthermore, exploring whether mammalian Piezo channels share analogous roles in density-dependent development may reveal deep evolutionary parallels in mechanobiology.

---

**Extended data Figure 1.**
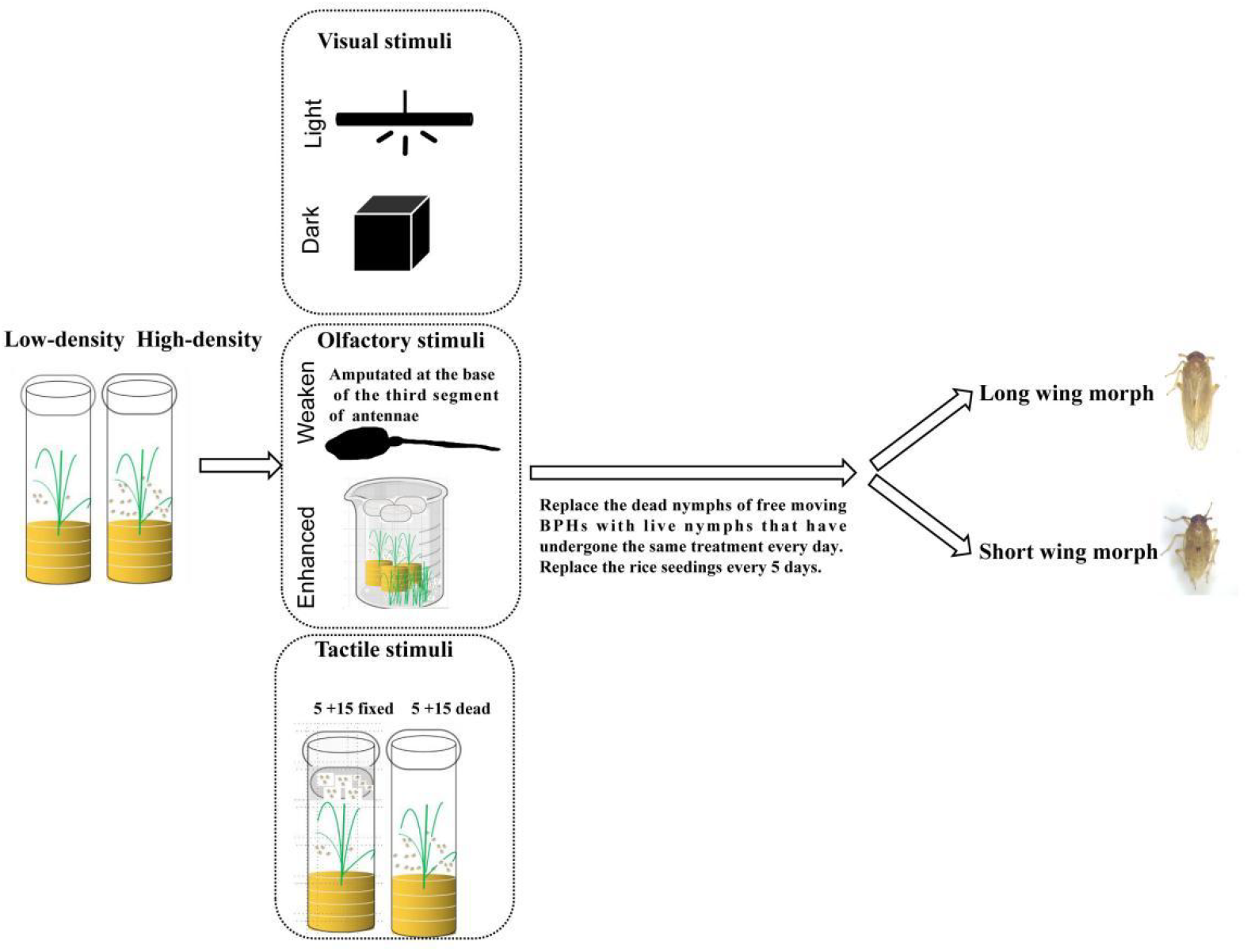
Experimental design to distinguish specific signaling modalities underlyin population density-induced wing dimorphism in *N*.*lugens*. Controls: BPHs were reared in low-density (5 nymphs/tube), or high-density (20 nymphs/tube) in regular light cycle (L:D=14:10) until emerged adults for all experiments control. Visual stimuli: BPHs were reared in low-density (5 nymphs/tube) or high-density (20 nymphs/tube) in regular light cycle light treatment or in 24 darkness until emerged adults, olfactory stimuli: it is hypothesized that olfactory was received via sensory organs at the base of the third segment of antennae. Weaken: to reduce olfactory sensation, antennae were amputated at the base of the third segment of antennae; Enhanced: to enhance phenomone concentration, 200 BPHs were kept outside the feeding vessel, but phenomone can pass through the vessels. tactile stimuli: 5 free-moving +15 dead: 5 BPHs were caged with the bodies of 15 dead BPHs, 5 free-moving +15 fixed: 5 BPHs were caged with 15 other live BPHs isolated by gauze fixation so that they were unable to touch. All treatments were reared in regular light cycle until emerged adults. Replace the dead nymphs of free moving BPHs with live ones which were parallel processed every day, and replace the rice seedlings every 5 days, then feeding until emerged adults.

**Extended data Figure 2.**
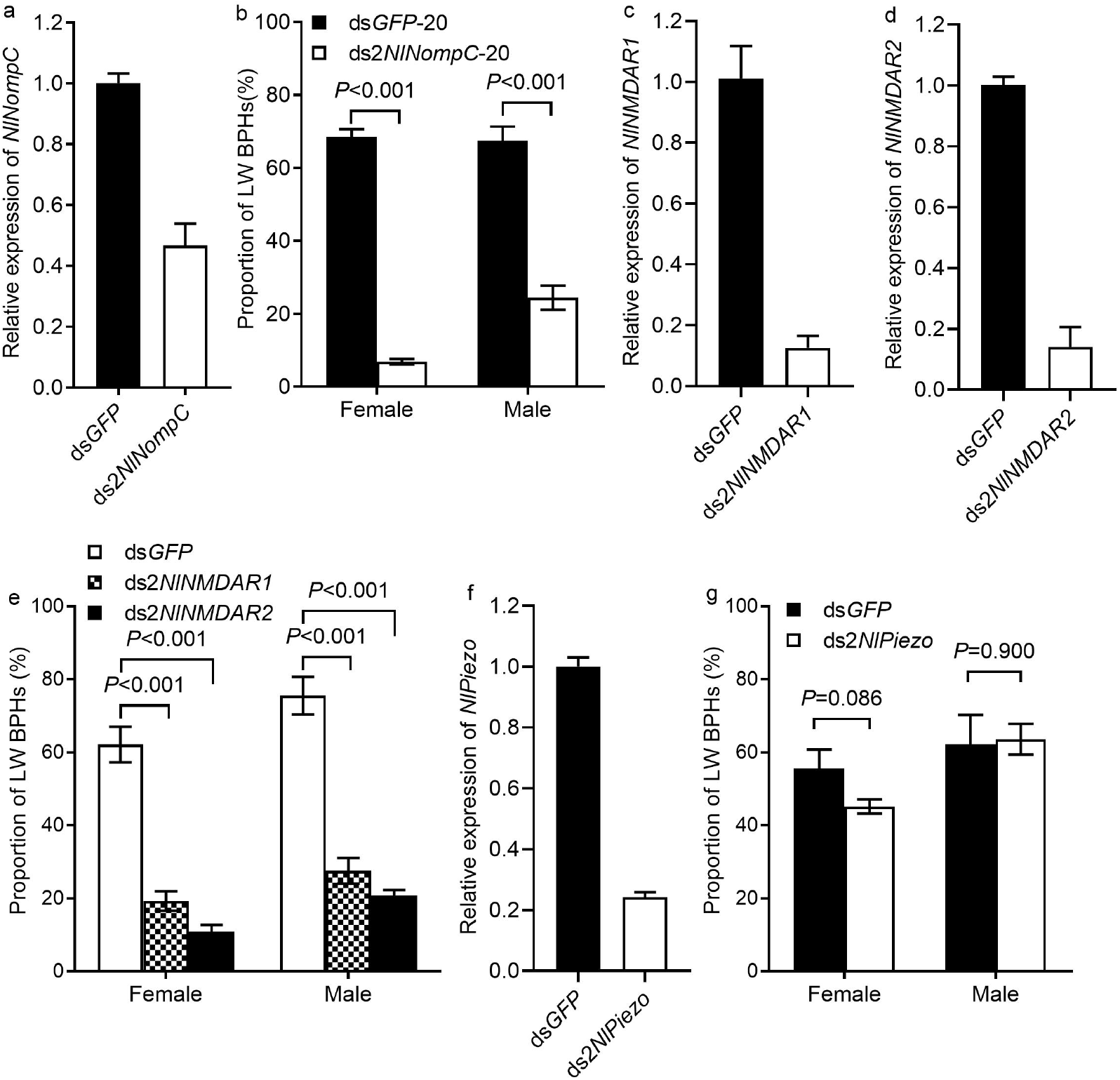
The 3rd-instar nymphs of BPHs were injected with dsRNA and reared in high density (20 nymphs/tube). a, Knockdown efficiency of the second dsRNA, ds2*NlNompC*, targeting *NlNompC*. b, The proportion of long-winged BPHs treated with ds*GFP* or ds2*NlNompC*, n=24-26. c, Knockdown efficiency of the second dsRNA, ds2*NlNMDAR1*, targeting *NlNMDAR1*. d. Knockdown efficiency of ds2*NlNMDAR2*, targeting *NlNMDAR2*. e. The proportion of long-winged BPHs ds*GFP* or ds2*NlNMDAR1/*ds2*NlNMDAR2*, n=19-30. f, Knockdown efficiency of the second dsRNA, ds2*NlPiezo*, targeting *NlPiezo*. g. The proportion of long-winged BPHs ds*GFP* or ds2*NlPiezo*. Mean ± s.e.m. from three independent experiments, n=12-22. (Independent samples *t*-test).

**Extended data Figure 3.**
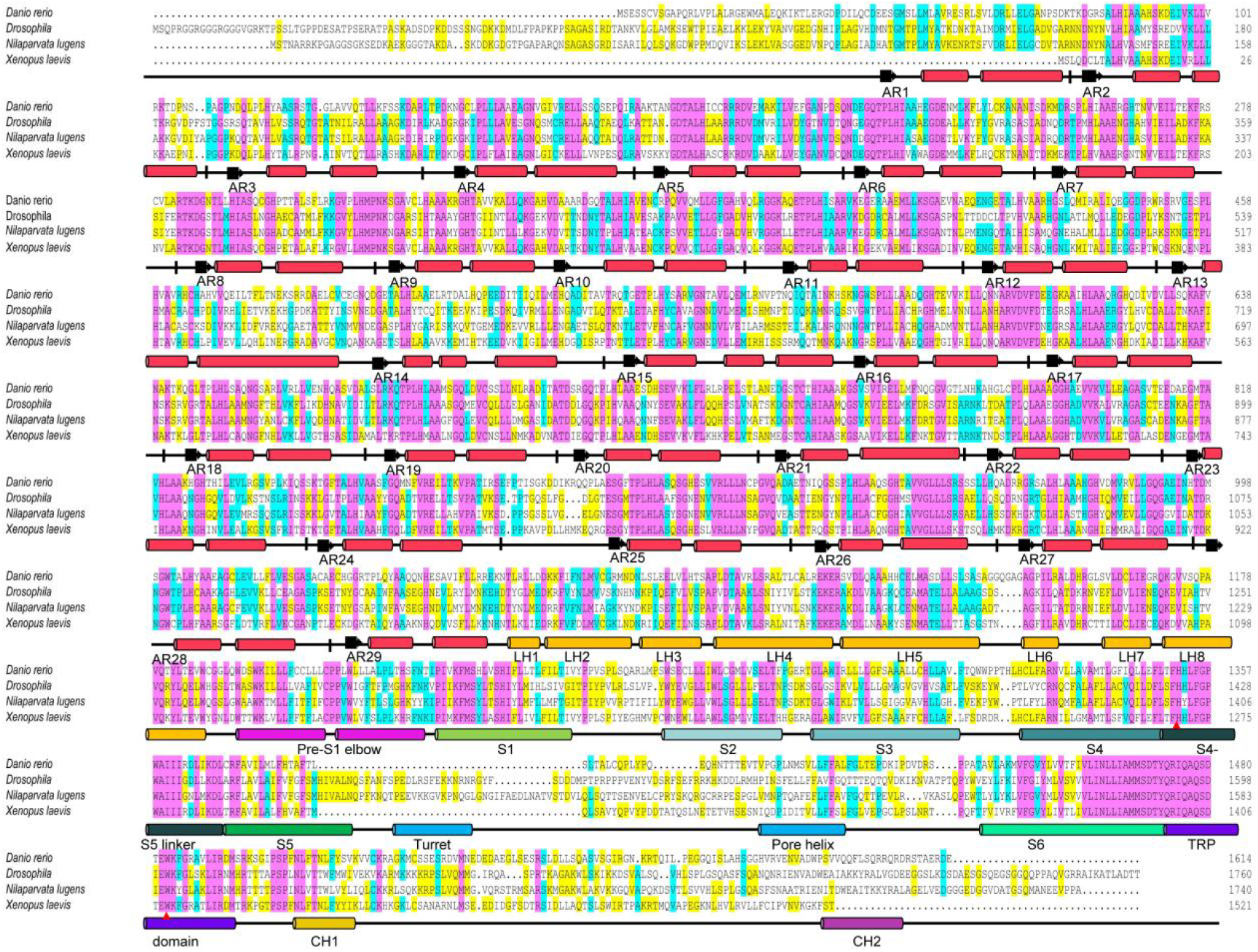
Sequence alignment of NompC orthologues. Sequence homology of NompC orthologues were analyzed by DNAMAN (83.13%). The conserved residues (the entire AR domain of 29 ARs, TRP domain and S4-S5 linker) are highlighted. Two critical amino acid residues (His1423 and Trp1601) in *Drosophila* NompC, essential for mechanogating (Jin et al., 2017), are evolutionarily conserved across orthologs and highlighted by red triangles in the alignment. Secondary structure elements are indicated above the sequence. *Xenopus laevis* (NP_001089174.1), *Danio rerio* (NP_899192.1), *Drosophila melanogaster* (NP_523483.2), *N. lugens* (XP_039285716.1).

**Extended data Figure 4.**
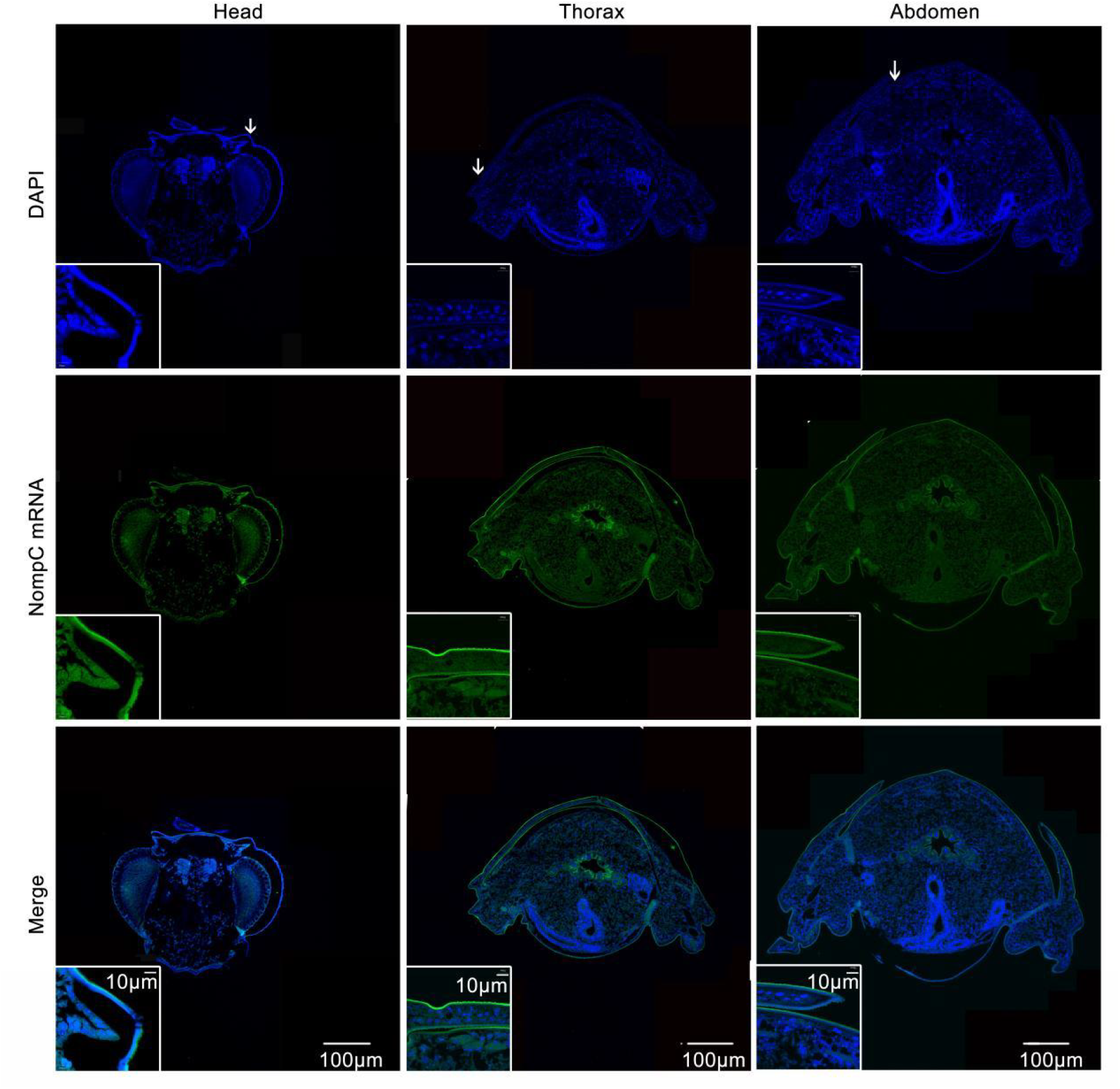
The expression of *NlNompC* mRNA was detected in the integument of the head, thorax, and abdomen using FISH. The oligonucleotide probe labeled with fluorochrome FAM (5’ primer sense) targets NlNompC mRNA expressed in epidermis. The arrow marks the area of local enlargement in the head, thorax, and abdomen, as shown at high magnification on the lower left. Scale bars: 100 μm.

**Extended data Figure 5.**
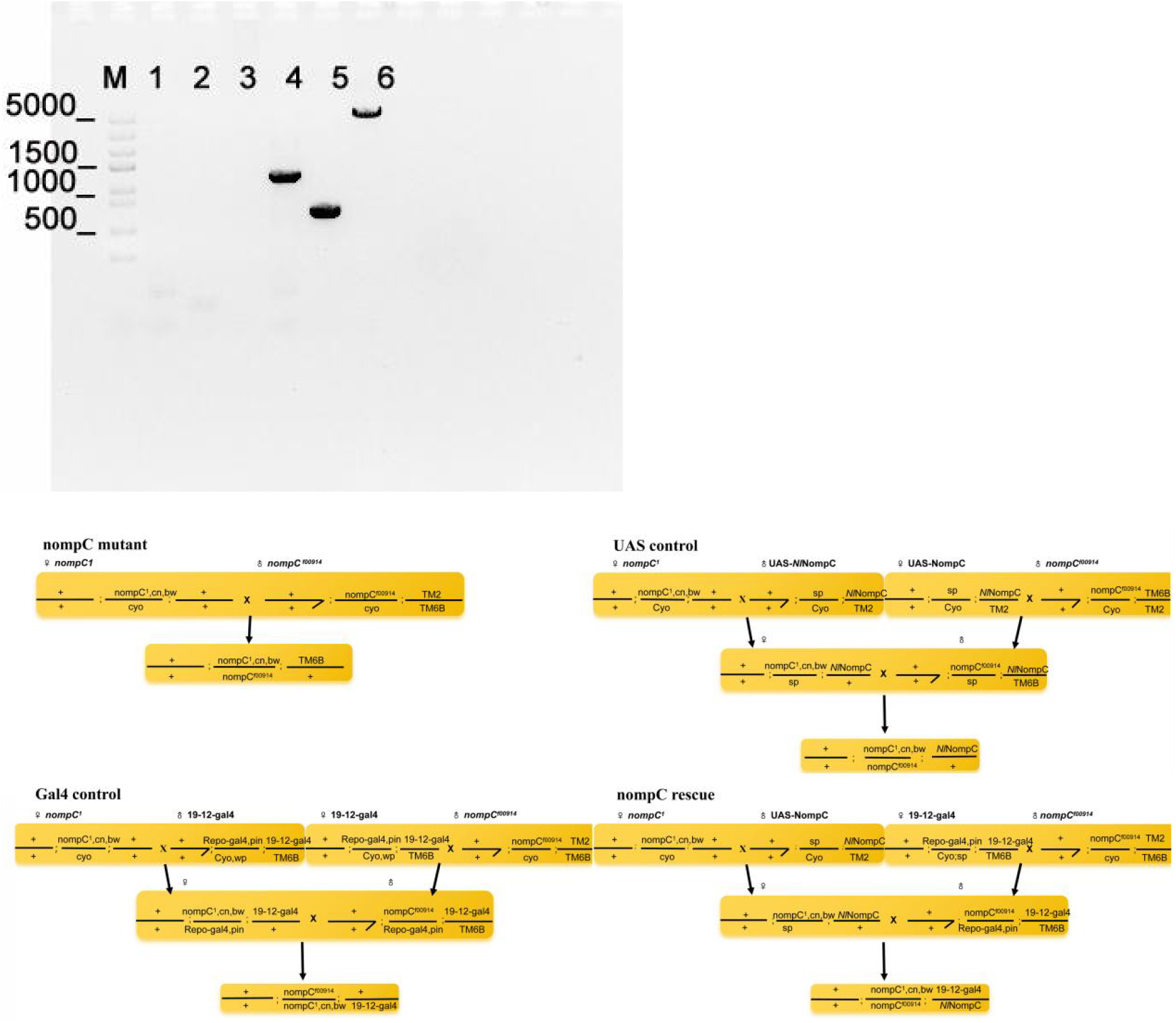
a. PUAST-NompC was inserted into the fruit fly strain (86F) detection. b. *Drosophila* hybridization scheme.

**Extended data Figure 6.**
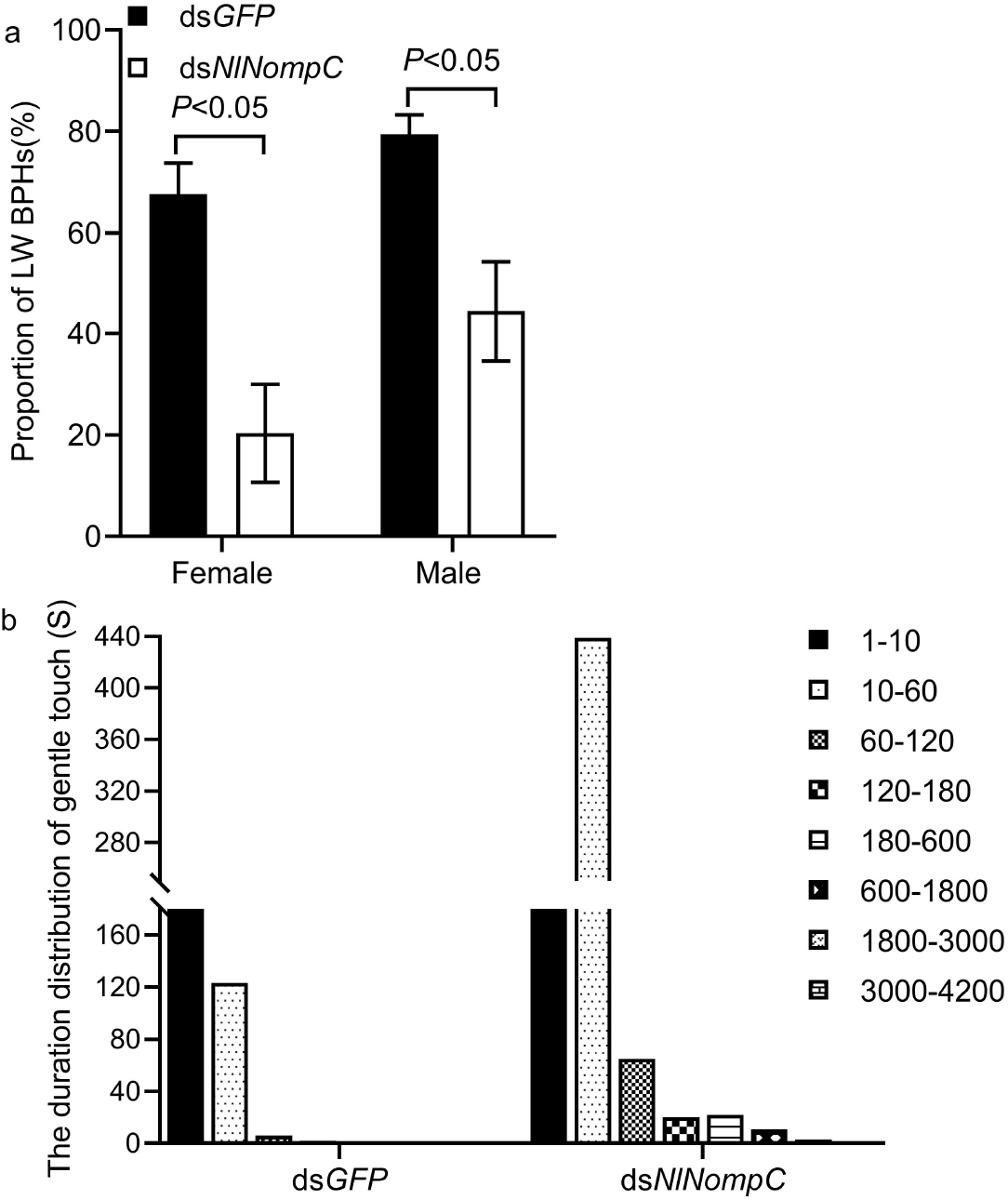
a. The wing-morph ratio of BPH in 3.5 ml container with density of 5 nymphs after treated with ds*GFP* or ds2*NlNompC* (n=20-22). b. The duration (seconds) of each gentile touch of planthoppers in 3.5 ml container with density of 5 nymphs treated with ds*GFP*, ds2*NlNompC* plus 2 uninjected nymphs of *S. furcifera*. n=22. (Independent samples *t*-test)

**Extended data Figure 7.**
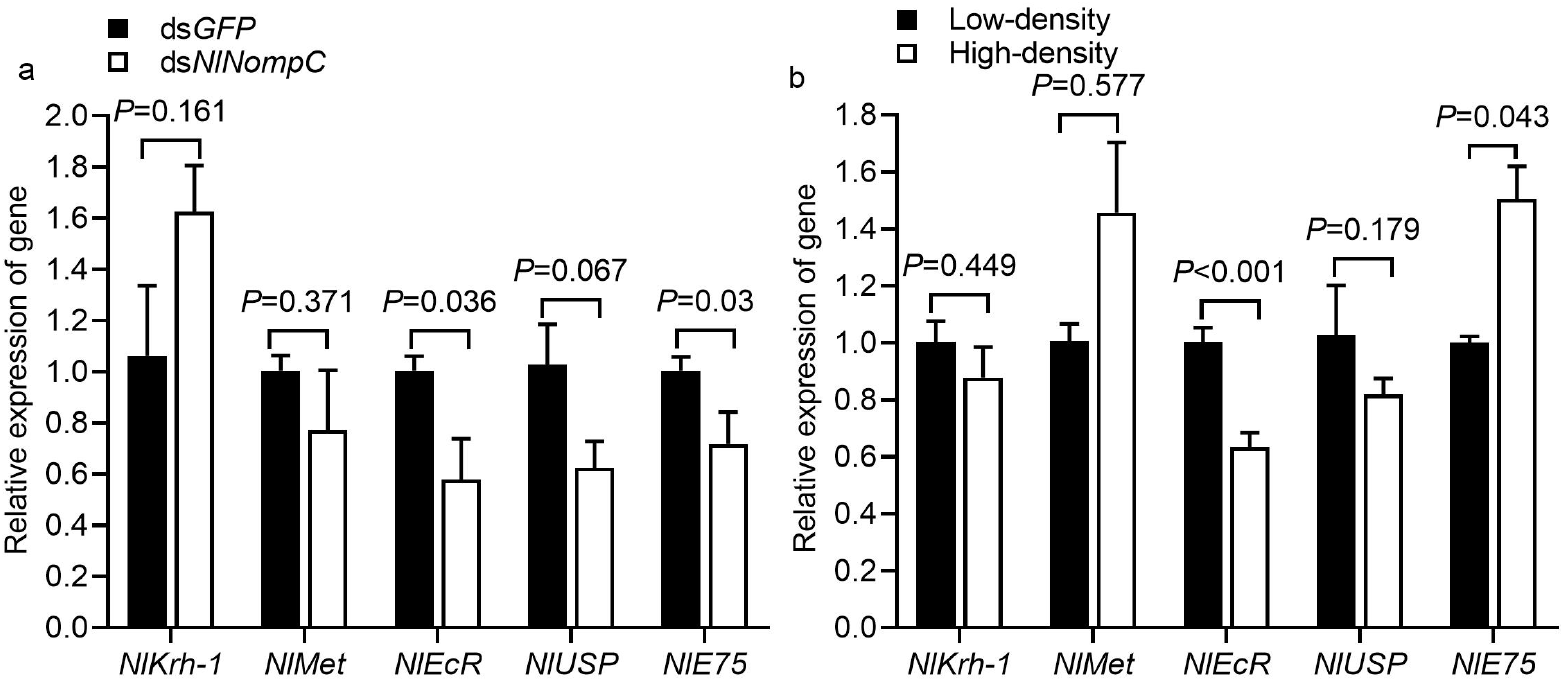
a. qPCR dection of the expression level of *NlKrh-1, NlMet, NlEcR, NlUSP*, and *NlE75* of 5th-instar nymphs after treated with ds*GFP* or ds*NlNompC*. b. qPCR dection of the expression level of *NlKrh-1, NlMet, NlEcR, NlUSP*, and *NlE75* of 5th-instar nymphs under low-density(5 nymphs/tube) or high-density (20 nymphs/tube) rearing conditions. n=4

**Extended data Figure 8.**
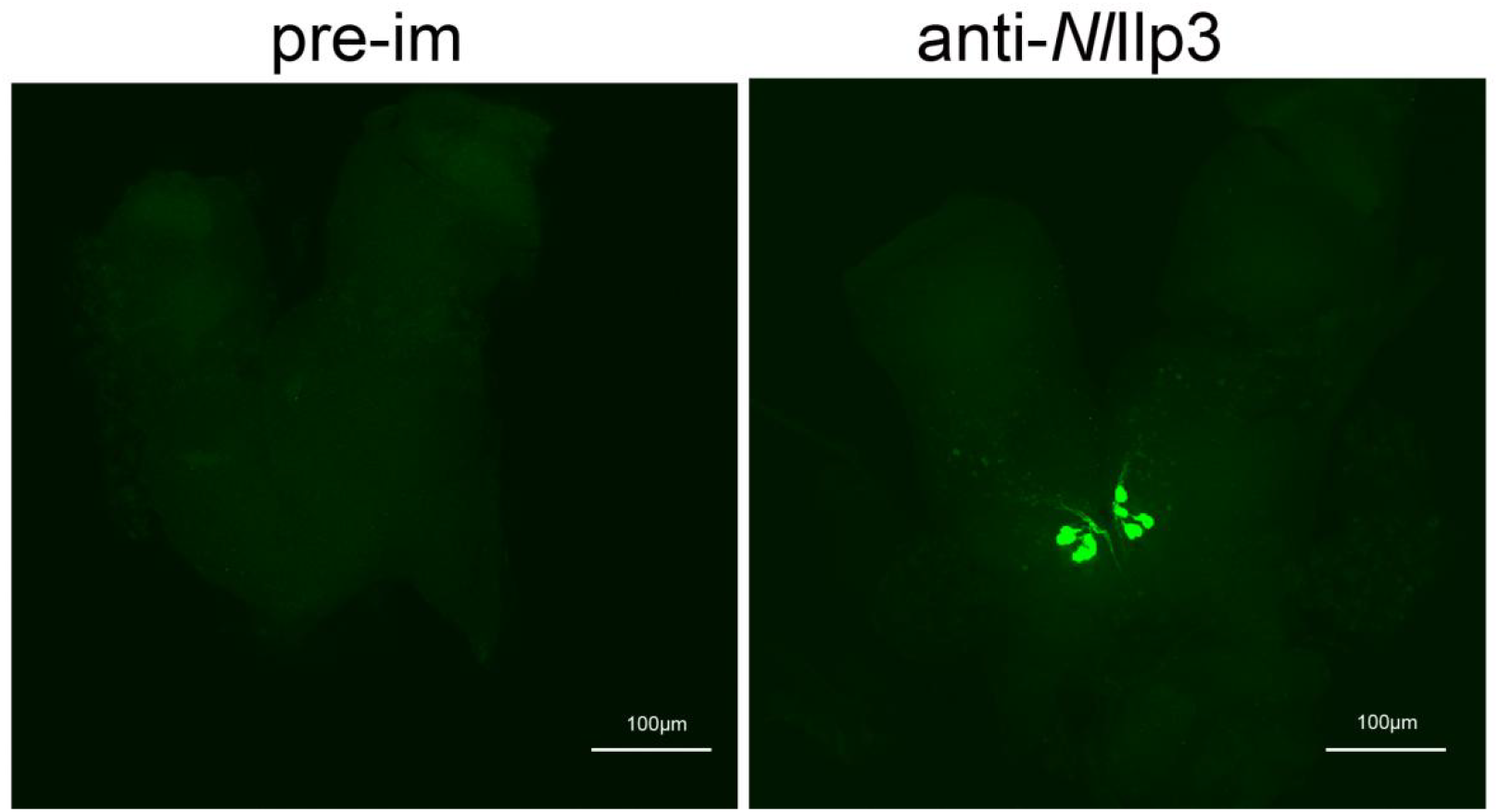
The specificity of anti-*Nl*Ilp3 antibodies was assessed by immunofluorescence with IgG purified from pre-immune serum (left) or anti-*Nl*Ilp3 (right).

**Extended data Figure 9.**
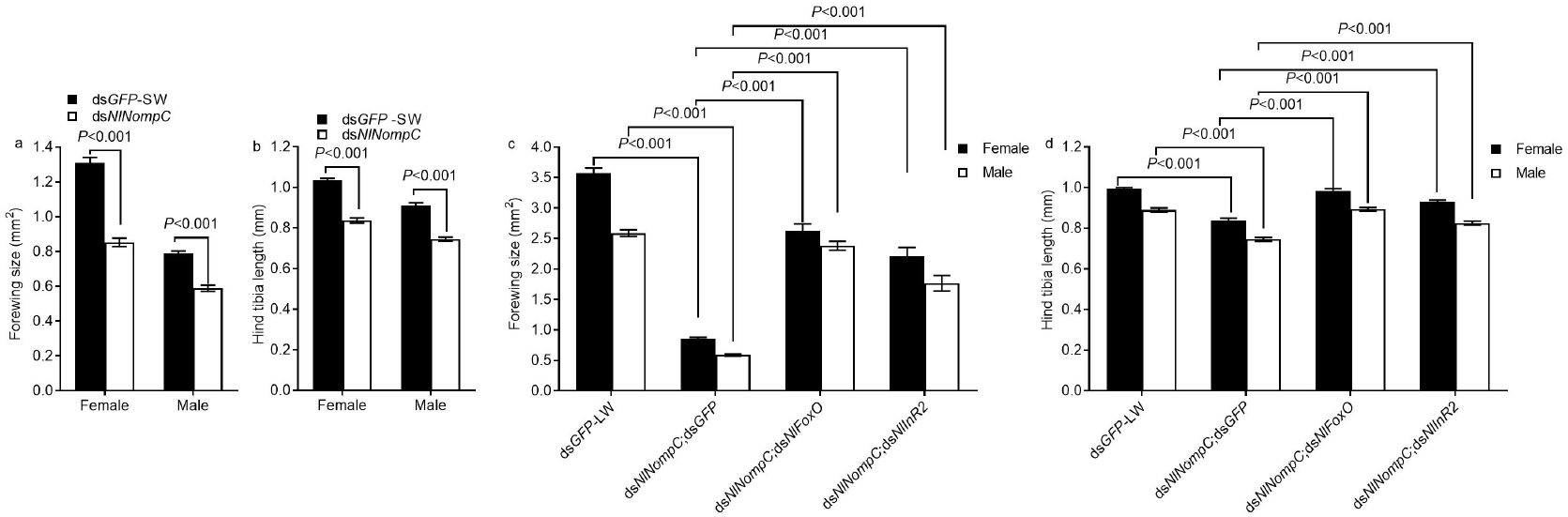
**a**, BPHs treated with ds*NlNompC* exhibited significantly reduced forewing size compared to ds*GFP*-SW controls. n=30-36. **b**, Similarly, hind tibia length was significantly reduced in ds*NlNompC*-treated BPHs relative to ds*GFP*-SW controls. n=30-36. c, Treatment of BPHs with ds*NlNompC*;ds*NlFox*O or ds*NlNompC*;ds*NlInR2* restored the forewing size phenotype observed in ds*NlNompC*-treated BPHs. n=24-36. d, Similarly, hind tibia length phenotype was restored in BPHs co-treated with ds*NlNompC*;ds*NlFox*O or ds*NlNompC*;ds*NlInR2* compared to ds*NlNompC*-treated controls. n=24-36. (Independent samples *t*-test)

**Extended data Figure 10.**
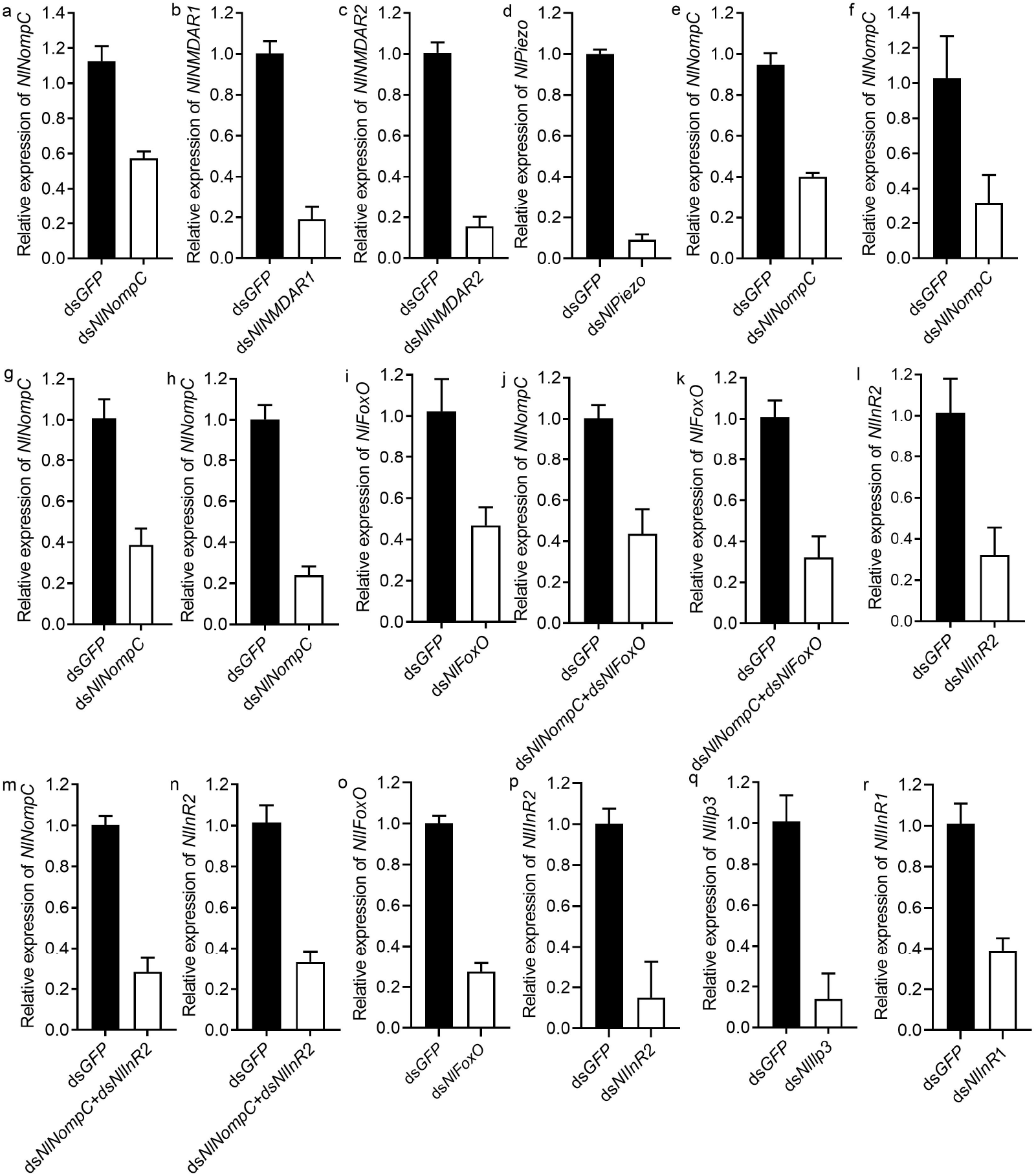
Examination of RNAi efficiency by qRT-PCR. a-r, Individual nymphs were pooled to extract total RNA after dsRNAs treatments, and cDNA was synthesized with random 8-mer oligo primers. The relative expression of each gene was normalized to the expression level of ribosomal actin. Mean ± s.e.m. from three experiments.

**Supplementary table S1**. The main primers used in this study.

**Movies S1-S4**. The Video recording of the gentle touch times and the duration of gentle touches among brown planthoppers in 3.5 ml container with density of 5 nymphs treated with ds*GFP* (Movies 1-2) or ds*NlNompC* (Movies 3-4)

**Movies S5-S8**. The Video recording of the gentle touch times and the duration of gentle touches among brown planthoppers in 3.5 ml container with density of 5 nymphs treated with ds*GFP* (Movies 5-6) or ds*NlNompC* (Movies 7-8) plus 2 uninjected nymphs of *S. furcifera*.

## Methods

### Insects

The *Nilaparvata lugens* (BPH), *Sogatella furcifera* (SFP) planthopper populations used in this work were originally collected in Huazhong Agricultural University experimental field of Wuhan, China, in 2021-2024, and density sensitivity of female BPHs with an increase of nymphal density. All insects were maintained in a growth chamber at 28 ^°^C (±1 ^°^C) under a photoperiod of 14 h :10 h (light: dark) at a relative humidity of 70% (± 5%) on rice seedlings (Taichuang Native 1). The rearing conditions were also applied to the following experiments.

### Density assay and the critical factor experiment of crowding-induced long-winged morph formation of BPH

For comparison of wing-from density relationships, 3th instar nymphs were put into test-vials (2 cm. in diameter and 9.5 cm. in height, 17ml) until hatching in the densities of 5, 10, 15, 20, 25, 30, with 4-5cm high of five rice plants in each for food. Each tube is filled with four absorbent sponges to provide water for the rice plants (2 cm. in diameter and 0.9 cm. in height). The rearing density of low-density (5 BPHs) and high-density BPHs (20 BPHs) were also applied to the following experiments.

The nymphs were divided into three groups and treated with visual, olfactory and tactile stimuli to study the most critical factors affecting long wing production. Visual stimuli: Light: low-density (5 BPHs), or high-density BPHs (20 BPHs) reared in regular photoperiod (14L:10D) until adult emergence; Dark: 5 BPHs, or 20 BPHs BPHs reared in darkness (in the box) until adult emergence; Olfactory stimuli: it is hypothesized that olfactory was received via sensory organs at the base of the third segment of antennae. 5 BPHs, or 20 BPHs reared in olfactory weakened conditions (antennae were amputated at the base of the third segment of antennae to reduce olfactory sensitivity) and olfactory enhanced conditions (200 BPHs (in 500 ml beaker) were kept outside of the feeding vials to increase olfactory sensitivity). Tactile stimuli: 5 free-moving +15 dead: 5 free-moving BPHs plus 15 dead BPHs, 5 free-moving +15 fixed: 5 free-moving BPHs plus 15 live BPHs wrapped with gauze so that two groups could not touch. Change food 5 days apart from all group density. Dead larvae were gently picked up from the feeding vials every day, replenish them to the corresponding density until hatching.

### Gentle touch times recording

We used an equal-scale reduction feeding vessels (4.5ml cuvettes, 1 cm ×1 cm ×4.5 cm 3.5ml) to observe the number of gentile touches by video under JT-H6 industrial microscope. The 3th instar larvae were put into test-vessels with 3-4cm high of one rice plants in each for food. Change food 2 days apart from all group density. Dead larvae were gently picked up from the vials every day, replenish them to the corresponding density until hatching. The gentle touch of each larva and the duration time per each gentle touch were recorded. The duration time per gentle touch was assessed with no response of larvae by touch. The number of gentle touch times indicates the average number of touches per larva. The duration time (s) per gentle touch classification level range is from 1 to 10, from 10 to 60, from 60 to 120, from 120 to 180, from 180 to 600, from 600 to 1800, from 1800 to 3000, from 3000 to 4200.

### dsRNA preparation and dsRNA microinjection in planthoppers

Synthesizing dsRNA and injection of planthoppers with dsRNA according to previously reported (Liu et al., 2020). The quantities of dsRNA injected into 3rd-instar nymphs were approximately 120 ng, respectively. The double dsRNA injection (90 ng/nymph for each dsRNA species) was injected into 3rd-instar nymphs. Each jar contained more than 600 individuals. Three days after the injections, RNA was extracted from 4 individuals to examine the gene silencing efficiency (Extended Data Fig. 10). Upon nymph molting, adult phenotypes were observed, and adult wing size and hind tibia length was measured using a stereomicroscope (Soptop szx12) with imageview. The primers designed in this study are shown in Supplementary Table 1.

### qRT–PCR analysis

Quantitative real-time PCR (qPCR) were performed following previously described methodologies (Tian *et al*., 2024). The primers designed in this study are shown in **Supplementary Table 1**.

### Pharmacological injection

FM1-43 reversibly blocks mechanotransduction (Yan et al 2013), MK-801 (dizocilpine) is a non-competitive NMDA receptor antagonist (Wong et al 1986, Geister et al 2008, Janus et al 2023) We injected the GdCl_3_ (6 × 10^−12^ µmol/nymph), MK-801 (6 × 10^−12^ µmol/nymph) or FM1-43 (1.2 × 10^−12^ µmol/nymph) to the third nymphs. Control group received the same volume (30µl) of ddH_2_O solution as that of the treated contorl solution. Then 20 nymphs were put into the plastic box and cultured together (2 cm. in diameter and 9.5 cm. in height, 17ml)).

### Sequence analysis

Sequence homology of NOMPC orthologues from X*enopus laevis, Danio rerio, Drosophila, Nilaparvata lugens* were analyzed by DNAMAN. The sequence similarity between *Drosophila* and *N*.*lugens* is 83.13%. The conserved residues (the entire AR domain of 29 ARs, TRP domain and S4-S5 linker) are highlighted according to the conserved of *Drosophila* NOMPC. The two residues (His1423 and Trp1572), which were shown in this study to be critical for mechanogating, are marked by red triangles. Secondary structure elements are indicated above the sequence (**Extended data Figure 3**).

### *In situ* hybridization

The oligonucleotide probe labeled with fluorochrome FAM (5’ primer sense) targeting *NlNompC*. In situ hybridization was carried out essentially as previously described (Liu et al 2020)

### Fly behavior assay

Fly stocks. The *NompC*^*1*^, Class III (*19-12-Gal4*) were from W. Zhang and NompC^f00914^ lines were isolated from Y. Zhu. The UAS-*Nl*NompC lines were generated in the lab. The behavioral assay for larvae’s response to gentle touch was carried out as described before (Yan et al., 2013). The larvae were allowed to crawl freely at room temperature. Flies were maintained under standard conditions at 25 °C. Embryo injections were carried out at Core technology facility of center for excellence in molecular cell science. CAS (http://http://sjzx.sibcb.ac.cn/Cn/Index/pageView/catid/108.html). The UAS-*Nl*nompC-attB gene expression vectors constructs were fixed point insertion at attP on the 3rd chromosome of 86F (Bloomington #24749).

### The gentle touch frequency recording

The JT-H6 industrial electron microscope (Shenzhen Jingtuo Youcheng) was used for continuous video recording for 24 hours to count the frequency of gentle touches and the contact time between nymphoses. After the video recording was completed, the rice seedlings were replaced every 2 days until the adult insects emerged, and the wing morph ratio was counted. The contact time starts to be counted when there is no response for more than 1 second. The duration (in seconds) of each gentle touch is classified into three levels: 1-10, 10-60, 60-120, 120-180, 180-600, 600-1800, 1800-3000, and 3000-4200.

### Preparation of His–*Nl*Ilp3 rabbit polyclonal serum

A PCR fragment (354 bp) encompassing the *NlIlp3* gene was amplified with the primers pIlp3-F and pIlp3-R (Supplementary Table 1), which were synthesized containing restriction endonuclease sites for NdeI and XhoI at their 5’ ends, respectively. The amplicons were cloned into the expression vector PET-28a with a 6 × His tag at the N terminus. The fusion proteins His–*Nl*Ilp3 were expressed in the *Escherichia colistrain* Rossetta (DE3) after induction with 0.5 mM isopropyl-b-D-thiogalactoside (IPTG) at 20^°^C for overnight. The samples were homogenized in buffer (8 M Urea,50 mM Tris, 300 mM NaCl, 0.1% Triton X-100, pH 8.0) and loaded onto a 5–12% SDS–PAGE gel (Beyotime). Polyclonal anti-serum against His–*Nl*Ilp3 was raised by immunizing rabbits with the purified proteins.

### Detection of *Nl*Ilp3 in brains

Heads of 5th-instar nymphs (0–24 h after ecdysis) were collected after treated with dsRNAs (120 ng per BPH) targeting *GFP, NlNompC* for 4 days and homogenized in RAPI (catalogue no. BL504A, Biosharp) containing protease inhibitor (catalogue no. P1262, Solarbio). Samples were quantified using the BCA method (catalogue no. P0012, Beyotime) and denatured in SDS–PAGE loading buffer. Equal amounts of protein (80 ug) were loaded for each lane on 15% SDS–PAGE gel. The b-actin polyclonal mouse serum (catalogue no. 2006068-8F10, ZENBIO) was used to monitor equal protein loading.

### Immunofluorescence assay

*Nl*Ilp3 antibody staining was carried out essentially as previously described (Tian et al., 2021). For *Nl*Ilp3 antibody staining, brains dissected from 5th-instar nymphs after treated with dsRNAs for 4 days, or from low-density and high-density rearing condition which fixed in 10% AF (95% ethanol absolute 90 ml, formalin 10 ml) for 6 h at room temperature. Then washed in PBT containing 0.2% Triton X-100, 0.1% Tween-20 four times for 15 min each. The specimens were transported in PBT for 1.5 h and then blocked in PBTA (0.2% Triton X-100, 0.1% Tween-20, 1.5% BSA) for 1 h, followed by incubation with primary rabbit serum against His–*Nl*Ilp3 (1:500) or GST–*Nl*NompC serum (1:500) in PBTA overnight at 4 ^°^C and then with goat anti-rabbit secondary antibody IgG/Alexa Flμor 488 (1:200) (catalogue no. D110061, Sangon Biotech) in PBTA for 3h after extensive washing. Finally washed in PBT six times for 15 min each. Fluorescence intensities were quantified using ImageJ software.

## Acknowledgements

H.X.H. was supported by the National Natural Science Foundation of China (32172396, 32372523). We thank Prof. Wei Zhang in School of Life Sciences for providing the *NompC*^*1*^ lines, 19-12-Gal4 lines and Prof. Yan Zhu in Chinese Academy of Sciences for providing the *NompC*^*f00914*^ lines.

## References

1. Liu F, Li X, Zhao M, Guo M, Han K, Dong X, Zhao J, Cai W, Zhang Q, Hua H. Ultrabithorax is a key regulator for the dimorphism of wings, a main cause for the outbreak of planthoppers in rice. Natl Sci Rev, 2020, 7(7):1181–1189.

2. Iwanaga K, Tojo S. Effects of juvenile hormone and rearing density on wing dimorphism and oocyte development in the brown planthopper, Nilaparvata lugens. J Insect Physiol, 1986, 32:585–590

3. Zhao J, Zhou Y, Li X, Cai W, Hua H. Silencing of juvenile hormone epoxide hydrolase gene (Nljheh) enhances short wing formation in a macropterous strain of the brown planthopper, Nilaparvata lugens. J Insect Physiol, 2017, 102:18–26

4. Ye XH, Xu L, Li X, He K, Hua HX, Cao ZH, Xu JD, Ye W, Zhang J, Yuan Z, Li F. miR-34 modulates wing polyphenism in planthopper. PLoS Genet, 2019, 15: e1008235

5. Simpson SJ, Despland E, Hagele BF, Dodgson T. Gregarious behavior in desert locusts is evoked by touching their back legs. PNAS USA, 2001, 98(7):3895–97.

6. Turner HN, Armengol K, Patel AA, Himmel NJ, Sullivan L, Iyer SC, Bhattacharya S, Iyer EPR, Landry C, Galko MJ, Cox DN. The TRP channels Pkd2, NompC, and Trpm act in cold-sensing neurons to mediate unique aversive behaviors to noxious cold in Drosophila. Curr Biol, 2016, 26(23):3116–3128

7. Wong EH, Kemp JA, Priestley T, Knight AR, Woodruff GN, and Iversen LL. The anticonvulsant MK-801 is a potent N-methyl-D-aspartate antagonist. PNAS, USA 1986, 83(18):7104–7108

8. Geister TL, Lorenz MW, Hoffmann KH and Fischer K. Effects of the NMDA receptor antagonist MK-801 on female reproduction and juvenile hormone biosynthesis in the cricket Gryllus bimaculatus and the butterfly Bicyclus anynana. J Exp Biol, 2008, 211: 1587–1593

9. Kim SE, Coste B, Chadha A, Cook B, Patapoutian A. The role of Drosophila Piezo in mechanical nociception. Nature, 2012, 483:209–212

10. Tracey WD Jr, Wilson RI, Laurent G, Benzer S. painless, a Drosophila gene essential for nociception. Cell, 2003, 18;113(2):261–73.

11. Li J, Zhang W, Guo Z, Wu S, Jan LY, Jan YN. A Defensive Kicking Behavior in Response to Mechanical Stimuli Mediated by Drosophila Wing Margin Bristles. J Neurosci, 2016, 2;36(44):11275–11282.

12. Jindra M, Palli SR, Riddiford LM. The juvenile hormone signaling pathway in insect development. Annu Rev Entomol, 2013, 58:181–204.

13. Yamanaka N, Rewitz KF, O’Connor MB. Ecdysone control of developmental transitions: lessons from Drosophila research. Annu Rev Entomol, 2013, 58:497–516.

14. Texada MJ, Koyama T, Rewitz K. Regulation of Body Size and Growth Control. Genetics, 2020, 216(2):269–313.

15. Kisimoto R. Effect of crowding during the larval period on the determination of the wing-form of an adult plant-hopper. Nature, 1956, 178:641–642

16. Saxena RC, Okech SH, Liquido NJ. Wing morphsim in the brown planthopper, Nilaparvata lugens. Insect Sci, 1981, 1:343–48

17. Iwanaga K, Nakasuji F, Tojo S. Wing polymorphism in Japanese and foreign strains of the brown planthopper, Nilaparvata lugens. Entomol Exp Appl, 1987, 43:3–10

18. Xu HJ, Xue J, Lu B, Zhang XC, Zhuo JC, He SF, Ma XF, Jiang YQ, Fan HW, Xu JY, Ye YX, Pan PL, Li Q, Bao YY, Nijhout HF, Zhang CX. Two insulin receptors determine alternative wing morphs in plan thoppers. Nature, 2015, 519:464–467

19. Zhang JL, Chen S J, Liu X Y. The transcription factor Zfh1 acts as a wing-morph switch in planthoppers. Nat commun, 2022, 13:5670

20. Zhang JL, Fu SJ, Chen SJ, Chen HH, Liu YL, Liu XY, Xu HJ. Vestigial mediates the effect of insulin signaling pathway on wing morph switching in planthoppers. PLoS Genet, 2021, 17(2):e1009312

21. Liu F, Li X, Zhao M, Guo M, Han K, Dong X, Zhao J, Cai W, Zhang Q, Hua H. Ultrabithorax is a key regulator for the dimorphism of wings, a main cause for the outbreak of planthoppers in rice. Natl Sci Rev, 2020, 7(7):1181–1189

22. Simpson SJ, Despland E, Hagele BF, Dodgson T. Gregarious behavior in desert locusts is evoked by touching their back legs. PNAS USA, 2001, 98(7):3895–97.

23. Ernst UR, Van Hiel MB, Depuydt G, Boerjan B, De Loof A, Schoofs L. Epigenetics and locust life phase transitions. J Exp Biol, 2015, 218(1):88–99

24. Johnson B. Wing polymorphism in aphids II. Interaction between aphids. Entomol Exp Appl, 1965, 8:49–64

25. Iwanaga K, Tojo S, Nagata T. Immigration of the brown planthopper, Nilaparvata lugens, exhibiting various responses to density in relation to wing morphism. Entomol Exp Appl, 1985, 38:101–18

26. Wang LX, Niu CD, Zhang Y, Jia YL, Zhang YJ, Zhang Y, Zhang YQ, Gao CF, Wu SF. The NompC channel regulates Nilaparvata lugens proprioception and gentle-touch response. Insect Biochem Mol Biol, 2019, 106:55–63

27. Cheng LE, Song W, Looger LL, Jan LY, and Jan YN. The role of the TRP channel NompC in Drosophila larval and adult locomotion. Neuron, 2010, 67:373–380.

28. Yan Z, Zhang W, He Y, Gorczyca D, Xiang Y, Cheng LE, Meltzer S, Jan LY, Jan YN. Drosophila NOMPC is a mechanotransduction channel subunit for gentle-touch sensation. Nature, 2013, 493:221–225

29. Guo XJ, Yu QQ, Chen DF, Wei JN, Yang PC, Yu J, Wang XH, and Kang L. 4-Vinylanisole is an aggregation pheromone in locusts. Nature, 2020, 584, 584–588

30. Zhang W, Cheng LE, Kittelmann M, Li J, Petkovic M, Cheng T, Jin P, Guo Z, Göpfert MC, Jan LY, Jan YN. Ankyrin repeats convey force to gate the NOMPC mechano transduction channel. Cell, 2015, 162(6):1391–1403

31. Yan C, Wang F, Peng Y, Williams CR, Jenkins B, Wildonger J, Kim HJ, Perr JB, Vaughan JC, Kern ME, Falvo MR, O’Brien 3rd ET, Superfine R, Tuthill JC, Xiang Y, Rogers SL, Parrish JZ. Microtubule acetylation is required for mechanosensation in Drosophila. Cell Rep, 2018, 25(4):1051–1065.e6

32. Rajendiran P, Jaafar F, Kar S, Sudhakumari C, Senthilkumaran B, Parhar IS. Sex Determination and Differentiation in Teleost: Roles of Genetics, Environment, and Brain. Biology (Basel), 2021, 10(10):973.

33. Ranade SS, Qiu Z, Woo SH, Hur SS, Murthy SE, Cahalan SM, Xu J, Mathur J, Bandell M, Coste B, Li YS, Chien S, Patapoutian A. Piezo1, a mechanically activated ion channel, is required for vascular development in mice. Proc Natl Acad Sci USA, 2014, 111(28):10347–52.

34. Li, J., Hou, B., Tumova, S. et al. Piezo1 integration of vascular architecture with physiological force. Nature, 2014, 515, 279–282.

35. Knell R. J. Population density and the evolution of male aggression. Journal of Zoology June 2009, 278(2):83–90.

36. Edwards PD, Frenette-Ling C, Palme R, Boonstra R. A mechanism for population self-regulation: Social density suppresses GnRH expression and reduces reproductivity in voles. J Anim Ecol, 2021, 90(4):784–795.

37. Rajendiran P, Jaafar F, Kar S, Sudhakumari C, Senthilkumaran B, Parhar IS. Sex Determination and Differentiation in Teleost: Roles of Genetics, Environment, and Brain. Biology (Basel), 2021, 10(10):973.

38. Guo X, Yu Q, Chen D, Wei J, Yang P, Yu J, Wang X, Kang L. 4-Vinylanisole is an aggregation pheromone in locusts. Nature, 2020, 584(7822):584–588.

39. Yang J, Yu Q, Yu J, Kang L, Guo X. 4-Vinylanisole promotes conspecific interaction and acquisition of gregarious behavior in the migratory locust. Proc Natl Acad Sci U S A, 2023, 120(37):e2306659120.

40. Lin XD, Gao H, Xu YL, Zhang YW, Li Y, Lavine MD, Lavine LC. Cell cycle progression determines wing morph in the polyphenic insect Nilaparvata lugens. iScience, 2020, 23(4): 2589–0042

41. Wong EH, Kemp JA, Priestley T, Knight AR, Woodruff GN, and Iversen LL. The anticonvulsant MK-801 is a potent N-methyl-D-aspartate antagonist. PNAS, USA 1986, 83(18):7104–7108

42. Geister TL, Lorenz MW, Hoffmann KH and Fischer K. Effects of the NMDA receptor antagonist MK-801 on female reproduction and juvenile hormone biosynthesis in the cricket Gryllus bimaculatus and the butterfly Bicyclus anynana. J Exp Biol, 2008, 211: 1587–1593.

43. Janus A, Lustyk K, Pytka K. MK-801 and cognitive functions: Investigating the behavioral effects of a non-competitive NMDA receptor antagonist. Psychopharmacology (Berl), 2023, 240(12):2435–2457.

